# Gut Microbiota-derived Acetate Safeguards the Colonic Epithelial Acetyl-CoA Reserve to Avert Colonic Senescence

**DOI:** 10.64898/2026.05.15.725523

**Authors:** Jie Du, Rajesh Sarkar, Lei Wang, Roya Amini Tabrizi, Lu Gao, Yan Li, Ashley Sidebottom, Hardik Shah, Mengjie Chen, Matthew Odenwald, Yan Chun Li

## Abstract

Aging is a major driver of tissue senescence, but little is known about the development of age-unrelated tissue senescence. Here we show that nucleocytosolic acetyl-CoA deficiency in colon epithelial cells, caused by *Acly* ablation and bacterial depletion, triggers age-independent, p53-dependent colonic senescence leading to severe systemic inflammation. Acetate supplementation, acetate-producing bacterial transplantation, targeted depletion of senescent cells or treatment with lysine deacetylase inhibitors blocks colonic senescence and inflammatory injury. Spontaneous colonic senescence develops following simultaneous deletion of epithelial *Acly* and *Acss2*, confirming that microbe-derived acetate maintains the epithelial acetyl-CoA pool via ACSS2 to avert colonic senescence. Mechanistically, acetyl-CoA deficiency deprives a cohort of mitochondrial and nuclear proteins of acetylation, leading to increased oxidative stress and DNA repair stress that trigger cellular senescence. Among these proteins, ATP5F1A-K161 acetylation and H4-K5/8 acetylation are required to protect colonic epithelial cells from developing senescence. These observations unveil a previously unknown mechanism that governs colonic senescence.

## Introduction

Cellular senescence is a state of irreversible cell cycle arrest associated with macromolecular damage and secretion of senescence-associated secretory phenotype (SASP), which includes cytokines, chemokines, growth factors and matrix remodeling proteases^1,2^. Senescent cells accumulate lysosomes that contain high levels of senescence-associated β-galactosidase (SA-β-gal), which is a well-accepted universal hallmark of senescence^3,4^. Senescence is induced in response to various stresses including telomere erosion, DNA damage, oxidative stress and oncogene activation^5,6^. These stress factors induce the activation of cell cycle inhibiting pathways including the p53/p21^Waf1/Cip1^ and p38/p16^Ink4a^ signaling pathways that converge to inhibit cyclin-dependent kinase (CDK)-cyclin complexes and activate retinoblastoma (Rb) protein to trigger cell cycle arrest and senescence^5,7–9^. SASP is a double-edged sword: it provides immune surveillance that recruits immune cells to clear senescent cells, which is an intrinsic design of the body to eliminate tumors or promote tissue remodeling and regeneration; however, persistent senescence or enduring SASP production that exceeds immune clearance can aggravate inflammatory injuries such as chronic inflammation, fibrosis, tissue dysfunction and cancer^5,10^.

Aging is a major driver of tissue senescence in animals and humans. It is believed that age-related accumulation of cellular senescence eventually leads to tissue senescence^1,5,11^, which is a major pathogenic factor responsible for the decline of tissue function and the increase in age-related tissue pathologies^7,12^. Like most tissues, the intestine exhibits age-related alterations and functional decline, but little is known about the molecular basis that drives intestinal senescence. One unique feature of the intestine is the persistent renewal of its epithelium every 4-5 days to maintain homeostasis. This continuous regeneration relies on the intestinal stem cells (ISCs)^13,14^, which replenish all differentiated cell types in the epithelium^15^. The age-related decline in the intestinal regenerative capacity is thought to result from alterations in the ISC niche and transcriptional factor expression^16–18^. The intestine is a complex system that involves constant interplays among the mucosal epithelial cells, the immune system and the microbiome^19,20^. There is evidence that gut microbiome influences gut senescence in the context of aging, and vice versa^8,21^, but exactly how gut microbiome regulates intestinal senescence remains unclear. Besides aging, tissue senescence is also driven by other factors; however, the causes of age-unrelated tissue senescence and its pathophysiological impacts remain poorly understood. In this regard, metabolic causes of senescence have been investigated in tissues such as the liver, pancreas and vasculature^22^, but little is known about metabolic control of intestinal senescence.

The constant regeneration of intestinal mucosal epithelium requires a synthesis of a great deal of biomass to meet the needs of cell division and differentiation. As acetyl-coenzyme A (acetyl-CoA) is the required substrate for the *de novo* lipogenesis and protein acetylation, we explored the role of acetyl-CoA in colon homeostasis. Our investigation leads to an unexpected discovery that epithelial nucleocytosolic acetyl-CoA deficiency triggers robust and widespread colonic senescence that causes severe local and systemic inflammation, and gut microbe-derived acetate helps maintain the epithelial acetyl-CoA pool to protect the host from developing colonic senescence. Moreover, our studies reveal that depletion of acetylation at specific lysine residues in H4 histone and mitochondrial ATP synthase is the key molecular event that kickstarts the senescence cascade triggered by acetyl-CoA deficiency.

## Results

### Critical roles of microbe-derived acetate in colon homeostasis

In mammalian cells citrate, an intermediary metabolite in the tricarboxylic acid (TCA) cycle, crosses mitochondrial membranes via transporter SLC25A1 to the cytosolic and nuclear compartments for the synthesis of acetyl-CoA catalyzed by ATP-citrate lyase (ACLY) (**Fig. S1A**)^23^. To understand the metabolic control of colonic mucosal homeostasis, we deleted *Acly* from colonic epithelial cells (CECs) by creating *Acly*^f/f^;*Cdx2*-Cre (CEC-*Acly*^−/−^) mice (**Fig. S1B,C**), as the *Cdx2* promoter drives Cre expression in epithelial cells from distal ileum to distal colon^24^. CEC-*Acly*^−/−^ mice were phenotypically normal including normal growth in both genders (**Fig. 1A, Fig. S1D**). To explain this result, we speculated that *Acly*^−/−^ epithelial cells may generate acetyl-CoA from gut bacteria-derived acetate to maintain their cellular functions (**Fig. S1A**). Gut bacteria produce short chain fatty acids (SCFAs), mainly including acetate, propionate and butyrate, from fermenting undigestible dietary fibers, and the microbe-derived SCFAs can enter colonic epithelial cells via membrane transporters SCL5A8 (SMCT1) and SLC16A1 (MCT1)^25–27^. Acetate can be converted to acetyl-CoA by acyl-CoA synthetase short chain family member 1 and 2 (ACSS1, ACSS2)^28^. ACSS1 is a mitochondrial enzyme^29,30^, whereas ACSS2 is a nucleocytosolic enzyme that converts cytosolic and nuclear acetate to acetyl-CoA^23,31^ (**Fig. S1A**). Epithelial *Acly* ablation had little effects on epithelial ACSS1 and ACSS2 expression (**Fig. S1B,C**). To test our hypothesis, we depleted gut bacteria by feeding mice an antibiotic cocktail (ABX) dissolved in the drinking water^32,33^ (**Fig. S1E,F**), and ABX had no effects on colonic ACLY and ACSS2 expression (**Fig. S1G**). Interestingly, all CEC-*Acly*^−/−^ mice died within two weeks following ABX (about 2-3 cycles of epithelial turnover), and supplementation of acetate, but not propionate or butyrate, in the drinking water was able to mostly rescue the mutant mice (**Fig. 1A,B**). As expected, ABX depleted fecal SCFAs including acetate, propionate and butyrate in both *Acly*^f/f^ and CEC-*Acly*^−/−^ mice, and acetate supplementation substantially raised fecal acetate levels in these mice (**Fig. 1C**). However, ABX with or without acetate supplementation had no effects on serum acetate levels (**Fig. 1D**), indicating that gut microbe-derived acetate contributes little to the circulating acetate. Consistent with our hypothesis, few organoids could be derived from CEC-*Acly*^−/−^ colonic crypts and the surviving *Acly*^−/−^ organoids grew poorly with few buddings, but supplementation of acetate, not propionate or butyrate, to the media rescued *Acly*^−/−^ organoids (**Fig. 1E,F**).

**Figure 1.**
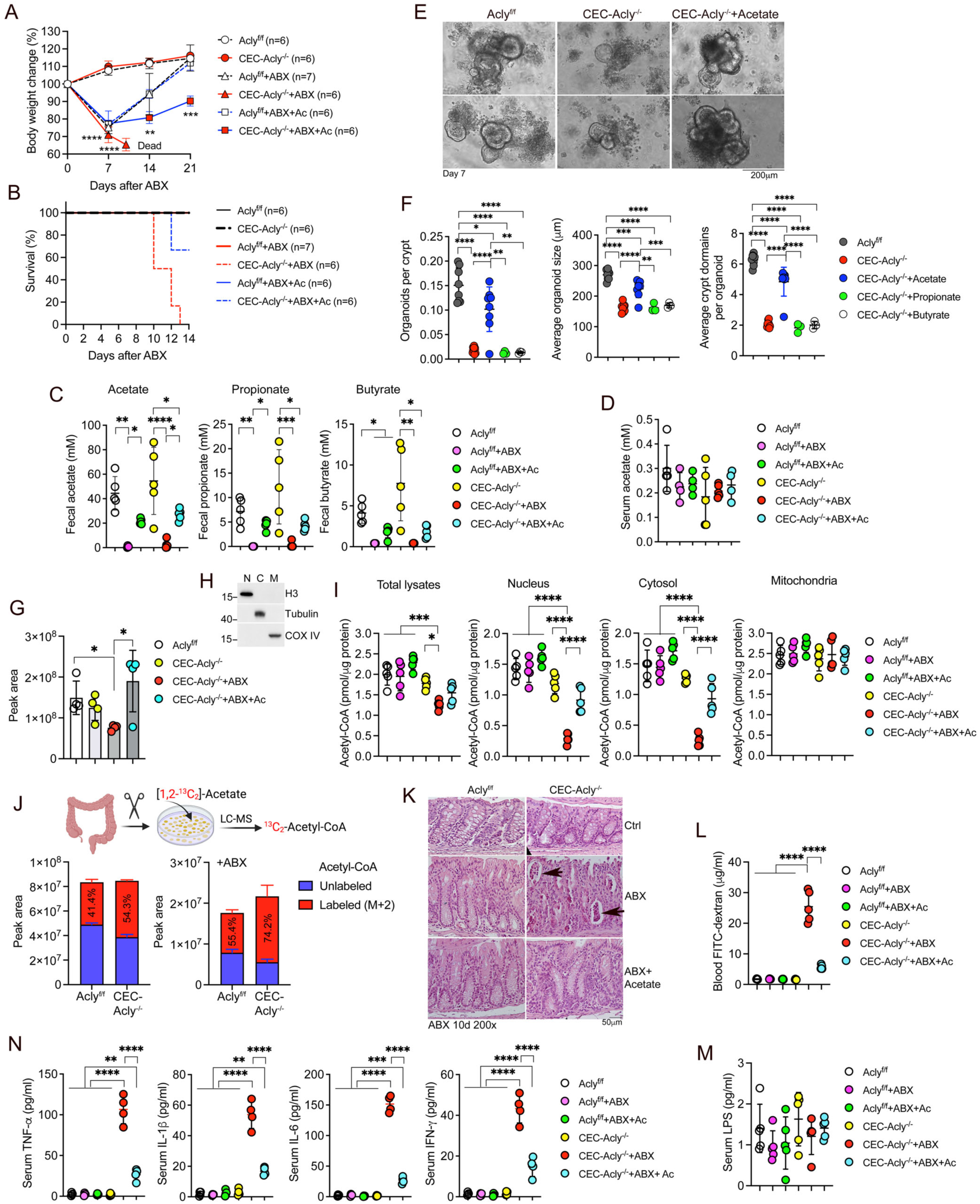
Gut microbiota-derived acetate contributes to the colonic epithelial acetyl-CoA pool to maintain colonic epithelial homeostasis. (A) Body weight changes in *Acly*^f/f^ and CEC-*Acly*^−/−^ mice following ABX and acetate (Ac) feeding in drinking water; **p<0.01, ***p<0.001, ****p<0.0001 vs. corresponding *Acly*^f/f^ controls, by Student’s t test; (B) Kaplan-Meier survival curves of *Acly*^f/f^ and CEC-*Acly*^−/−^ mice following ABX and Ac feeding; (C) Fecal acetate, propionate and butyrate concentrations in *Acly*^f/f^ and CEC-*Acly*^−/−^ mice on day 10 post ABX and Ac feeding, n=5 each group; (D) Serum acetate concentrations in *Acly*^f/f^ and CEC-*Acly*^−/−^ mice on day 10 following ABX and Ac feeding; (E) Photograph images of day 7 colonic organoids derived from *Acly*^f/f^ and CEC-*Acly*^−/−^ mice with or without acetate supplementation in the media; (F) Quantitation of organoid forming efficiency, organoid size and crypt domains per organoid in *Acly*^f/f^ and CEC-*Acly*^−/−^ colonic organoids cultured in the presence of acetate, propionate or butyrate in the media; (G) Mass spectrometric quantitation of acetyl-CoA levels in colonic crypt lysates prepared from *Acly*^f/f^and CEC-*Acly*^−/−^ mice on day 7 after ABX and Ac feeding; (H) Western blot analysis to confirm the purity of nuclear (N), cytosolic (C) and mitochondrial (M) fractions prepared from purified colonic crypt cells; (I) Quantitation of acetyl-CoA concentration in total lysates and nuclear, cytosolic and mitochondrial fractions of purified colonic crypts from *Acly*^f/f^ and CEC-*Acly*^−/−^ mice on day 7 following ABX and acetate feeding; (J) Quantitation of ^13^C_2_-acetyl-CoA generation within 60 min from [1,2-^13^C]-acetate in colonic tissue cultures derived from *Acly*^f/f^ and CEC-*Acly*^−/−^ mice with or without ABX for 7 days; n=4 each group; (K) H&E stained sections of distal colon from *Acly*^f/f^ and CEC-*Acly*^−/−^ mice on day 10 after ABX and acetate feeding; *Arrows* indicate detached crypts; (L) Blood FITC-dextran concentrations measured at 2.5 hours after FITC-dextran gavage in *Acly*^f/f^and CEC-*Acly*^−/−^ mice on day 10 after ABX and acetate feeding; n=5 each group; (M) Serum LPS concentrations in *Acly*^f/f^ and CEC-*Acly*^−/−^ mice on day 10 after ABX and acetate feeding; (N) Serum TNF-α, IL-1β, IL-6 and IFN-γ concentrations in *Acly*^f/f^ and CEC-*Acly*^−/−^ mice on day 10 after ABX and acetate feeding; n=5 each group. Data are presented as mean ± SD. *p<0.05; **p<0.01; ***p<0.001, ****p<0.0001 by two-way ANOVA.

We then quantified colonic epithelial acetyl-CoA concentrations by mass spectrometry. Metabolomic analysis confirmed that bacterial depletion significantly reduced acetyl-CoA levels in colonic epithelial cells of CEC-*Acly*^−/−^ mice, which was reversed by acetate feeding (**Fig. 1G**). This explains why acetate feeding is able to rescue ABX-treated CEC-*Acly*^−/−^ mice. Further analysis of subcellular fractions revealed a dramatic reduction of acetyl-CoA only in the nuclear and cytosolic compartments, not in mitochondria, in ABX-treated CEC-*Acly*^−/−^ colonic epithelial cells, and acetate feeding substantially rescued the nuclear and cytosolic acetyl-CoA deficiency (**Fig. 1H,I**). We further performed *ex vivo* [1,2-^13^C]-acetate tracing in colon organ cultures, which confirmed the formation of ^13^C_2_-acetyl-CoA in CEC-*Acly*^−/−^ colonic cells, but less in *Acly*^f/f^ colonic cells, with or without ABX (**Fig. 1J, Fig. S1H,I**). The global cellular metabolic profiling showed that epithelial *Acly* depletion, with or without ABX and with or without acetate supplementation, had no significant impacts on most of the intermediary metabolites related to the TCA cycle, except for α-ketoglutarate, succinate, glutamate and 2-hydroxyglutarate (**Fig. S2**), suggesting that the mitochondrial TCA cycle remains relatively intact in these colonic epithelial cells.

One important question is why all CEC-*Acly*^−/−^ mice died within two weeks following ABX? Histological examinations revealed an abnormal colonic epithelial morphology with severe depletion of mucus-producing goblet cells in ABX-treated CEC-*Acly*^−/−^ mice (**Fig. S1J**). High power magnification showed a collapsed crypt structure in the colonic mucosa, where the crypts were detached *en masse* from the basement membrane and shrank into a packed cell mass (**Fig. 1K**). As a result, mucosal permeability was markedly increased in CEC-*Acly*^−/−^+ABX mice (**Fig. 1L**), but their circulating LPS levels were not escalated because of microbial depletion (**Fig. 1M**). Unexpectedly, circulating inflammatory cytokines (TNF-α, IL-6, IL-1β, IFN-γ) were dramatically elevated in ABX-treated CEC-*Acly*^−/−^ mice (**Fig. 1N**), leading to severe lung and liver injuries (**Fig. S1K,L,M**). Apparently, this “sterile” systemic inflammation is not caused by bacterial invasion, and the multi-organ injury caused by the cytokine storm-like systemic inflammation is the cause of death for CEC-*Acly*^−/−^+ABX mice. As expected, acetate supplementation mostly normalized the colonic epithelial structure (**Fig. 1K, Fig. S1J**), corrected the mucosal permeability (**Fig. 1L**), lowered the circulating cytokine levels (**Fig. 1N**), and prevented the organ injury (**Fig. S1K,L,M**) in CEC-*Acly*^−/−^+ABX mice. Collectively, these observations demonstrate that gut microbe-derived acetate contributes to colonic epithelial acetyl-CoA synthesis and plays a critical role in the maintenance of colonic homeostasis.

### Acetyl-CoA deficiency triggers robust age-unrelated, p53-dependent colonic senescence

In a leaky gut an increase in microbiota invasion is usually a cause of systemic inflammation; surprisingly, however, the depletion of intestinal bacteria could still trigger a severe systemic inflammation in CEC-*Acly*^−/−^ mice. To explore the underlying mechanism, we examined the transcriptomic profiles of colonic epithelial cells isolated from *Acly*^f/f^, CEC-*Acly*^−/−^, CEC-*Acly*^−/−^+ABX and CEC-*Acly*^−/−^+ABX+Acetate (Ac) mice by bulk RNA-seq (**Fig. S3A**). Only 373 differentially expressed genes (DEGs) were detected between *Acly*^f/f^ and CEC-*Acly*^−/−^ mice (**Fig. S3B**), but ABX markedly increased DEGs between *Acly*^f/f^ and CEC-*Acly*^−/−^+ABX mice (2688) (**Fig. S3C**) and between CEC-*Acly*^−/−^ and CEC-*Acly*^−/−^+ABX mice (2239) (**Fig. S3E**), and large numbers of DEGs remained seen following acetate supplementation (2477 between *Acly*^+/+^ and CEC-*Acly*^−/−^+ABX+Ac; 3117 between CEC-*Acly*^−/−^ and CEC-*Acly*^−/−^+ABX+Ac; 1669 between CEC-*Acly*^−/−^+ABX and CEC-*Acly*^−/−^+ABX+Ac) (**Fig. S3D,F,G**), suggesting that acetate feeding was not able to normalize the transcriptional effect of ABX. Reactome pathway analysis revealed that the most predominant pathways that were activated are related to mitotic prophase chromosome condensation, nucleosome assembly, telomere maintenance, DNA double strand break (DSB) response, DNA damage response (DDR), DDR/telomere stress-induced senescence and cellular senescence in the CEC-*Acly*^−/−^ colons, and the enrichment of these pathways was further enhanced by ABX, but mostly normalized by acetate feeding (**Fig. 2A**). These data suggest that the DNA repair machinery ramps up to maintain a homeostasis in the absence of citrate to acetyl-CoA conversion, and maximizes its capacity in severe acetyl-CoA deficiency when the homeostasis is broken down. DNA repair stress or DDR is a major inducer of senescence^3,34^. A large number of histone variants were induced in the CEC-*Acly*^−/−^colons, which may reflect the fact that ACLY is involved in DNA repair^35^, and this induction was enhanced further by ABX but normalized by acetate feeding (**Fig. 2B**). On the other hand, the expression of centromere proteins and cyclins (*Ccna2, Ccnb1, Ccnb2, Ccnd1*) was markedly suppressed in the CEC-*Acly*^−/−^ and CEC-*Acly*^−/−^+ABX colons, whereas cyclin dependent kinase (CDK) inhibitors including a well-accepted senescence hallmark *Cdkn1a* (p21^Waf1/Cip1^) were highly induced by ABX in the CEC-*Acly*^−/−^+ABX colons and partially normalized by acetate supplementation (**Fig. 2C**). These observations revealed the existence of DNA repair stress, inhibition of chromosome segregation in mitosis, cell cycle arrest and development of colonic senescence in ABX-treated CEC-*Acly*^−/−^ mice.

**Figure 2.**
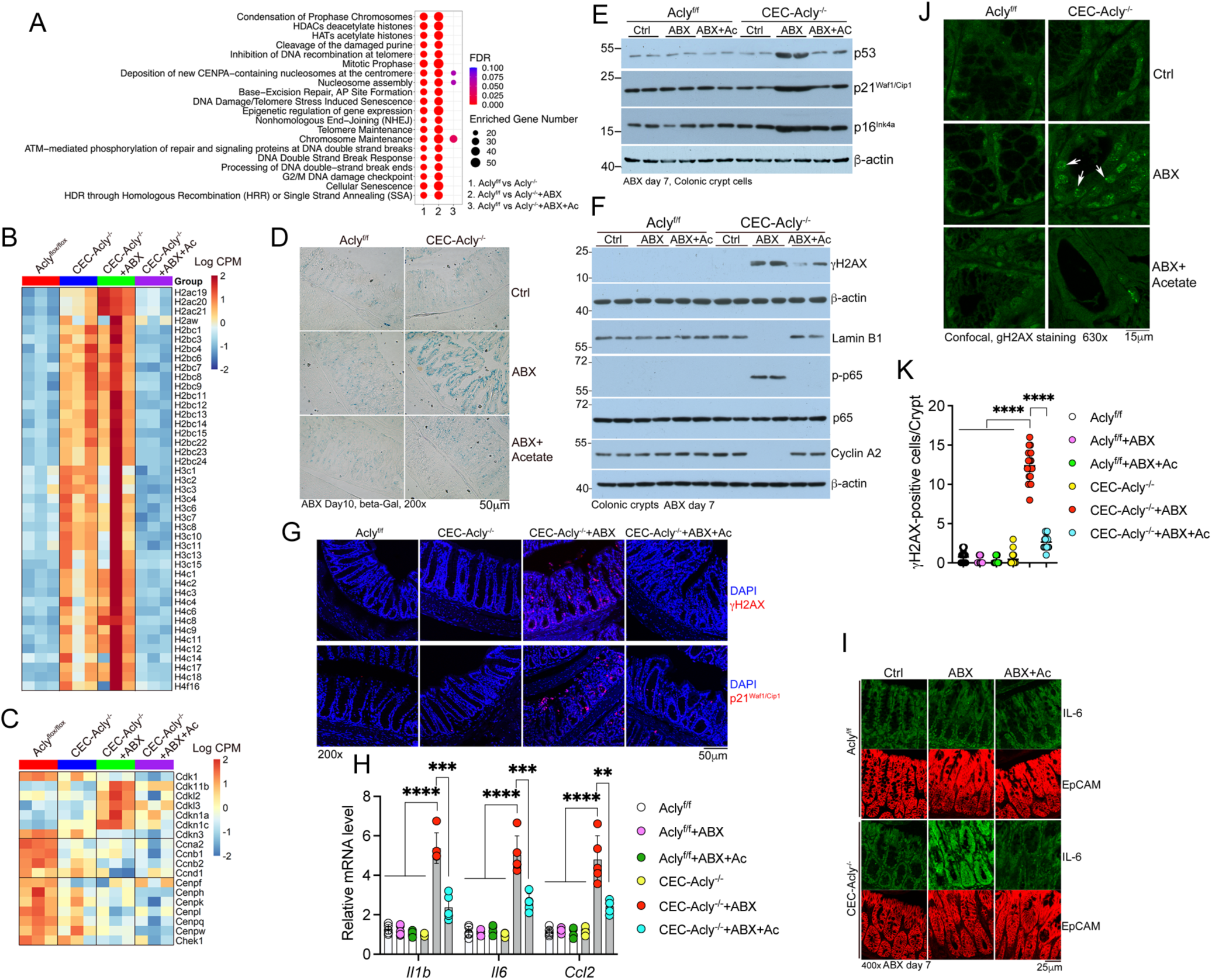
Epithelial acetyl-CoA deficiency triggers robust p53-dependent colonic senescence. (A) Top Reactome pathways identified in the differentially expressed genes (DEGs) between *Acly*^f/f^ and CEC-*Acly*^−/−^ colonic crypt cells with or without ABX and acetate (Ac) treatment for 5 days; (B) Heatmap showing changes in the expression of histone variants in colonic crypt cells from *Acly*^f/f^ and CEC-*Acly*^−/−^ mice with or without ABX and Ac treatment for 5 days; (C) Heatmap showing changes in the expression of centromere protein genes and genes involved in cell cycle regulation in colonic crypt cells from *Acly*^f/f^ and CEC-*Acly*^−/−^ mice with or without ABX and Ac treatment for 5 days; (D) Senescence-associated β-galactosidase (SA-β-gal) staining in frozen colon sections from *Acly*^f/f^ and CEC-*Acly*^−/−^ mice on day 10 following ABX and Ac feeding; (E) Western blot assessment of canonical senescence hallmarks in colonic crypt cells from *Acly*^f/f^ and CEC-*Acly*^−/−^ mice on day 7 following ABX and Ac treatment; (F) Western blot assessment of senescence-related markers in colonic crypt cells from *Acly*^f/f^ and CEC-*Acly*^−/−^ mice on day 7 following ABX and Ac treatment; (G) Immunofluorescence staining of γH2AX and p21^Waf1/Cip1^ in colonic sections from *Acly*^f/f^ and CEC-*Acly*^−/−^ mice on day 7 following ABX and Ac treatment; (H) RT-qPCR quantitation of SASP cytokines in colonic crypt cells from *Acly*^f/f^ and CEC-*Acly*^−/−^ mice on day 7 following ABX and Ac treatment; (I) Immunofluorescence staining of IL-6 and EpCAM in colonic sections from *Acly*^f/f^ and CEC-*Acly*^−/−^mice on day 7 following ABX and Ac treatment; (J) Confocal microscopic images of γH2AX immunofluorescence staining in colonic sections from *Acly*^f/f^ and CEC-*Acly*^−/−^ mice on day 7 following ABX and Ac treatment. *Arrows* indicate γH2AX loci in the nucleus of crypt epithelial cells; (K) Quantitation of γH2AX-positive cells per mucosal crypt in colonic sections from *Acly*^f/f^ and CEC-*Acly*^−/−^ mice on day 7 following ABX and Ac treatment; Data are presented as mean ± SD. *p<0.05; **p<0.01; ***p<0.001, ****p<0.0001 by two-way ANOVA.

We further performed analyses to confirm the development of colonic senescence. As expected, the CEC-*Acly*^−/−^ colonic crypts exhibited strong SA-β-gal staining following ABX (**Fig. 2D**), and Western blot and immunostaining analyses showed a robust induction of canonical hallmarks of senescence (p53, p21^Waf1/Cip1^ and p16^Ink4a^ ^3^) and suppression of cyclin A2 in these mice (**Fig. 2E,F,G**). Moreover, local SASP (*Il1b, Il6,* and *Ccl2*), another senescence hallmark ^1,2^, was also highly induced by ABX in CEC-*Acly*^−/−^ colonic mucosa (**Fig. 2H**). Co-immunostaining using antibodies against EpCAM (epithelial marker) and IL-6 (the most common SASP) confirmed that the robust SASP production was from colonic epithelial cells in ABX-treated CEC-*Acly*^−/−^ mice (**Fig. 2I**). Given the large surface area of the colonic epithelium, the amount of SASP continuously secreted from the senescent cells was sufficient to trigger a cytokine storm in the circulation (see **Fig. 1N**), causing multi-organ injuries and premature death. Acetate feeding substantially alleviated these senescence phenotypes in ABX-treated CEC-*Acly*^−/−^ mice (**Fig. 2D-I**).

DSBs cause accumulation of cytoplasmic chromatin fragments (CCFs) that activate the cGAS-STING-NF-κB pathway to induce SASP^34,36–38^. Presence of γH2AX loci or induction of γH2AX is an early response to DSBs and a hallmark of DNA damage^3,34,39^. Loss of Lamin B1, another biomarker of senescence, compromises nuclear integrity leading to CCF appearance^3,40^. We found that γH2AX loci were markedly increased in colonic epithelial cells of CEC-*Acly*^−/−^ mice following ABX (**Fig. 2J,K**), and γH2AX protein and NF-κB p65 phosphorylation (p-p65) were highly induced, whereas Lamin B1 was drastically reduced in ABX-treated CEC-*Acly*^−/−^ colonic crypts (**Fig. 2F,G**); all these changes were reversed or corrected following acetate supplementation (**Fig. 2F,G,J,K**).

It is well known that p53 is required for the development of senescence^3,9,37^. Activation of p53 in response to DNA damage induces p21^Waf1/Cip1^ that triggers cell cycle arrest and senescence^9^. To address whether p53 is required for the development of colonic senescence, we generated *Acly^f/f^*;Cdx2-Cre;*Trp53*^−/−^ (CEC-*Acly^−/−^*;*Trp53*^−/−^) mice that were depleted of both ACLY and p53 from colonic epithelial cells (**Fig. S4A,B**). CEC-*Acly*^−/−^;*Trp53*^−/−^ mice showed normal growth with no obvious abnormalities (**Fig. S4C**); however, following ABX treatment, in contrast to CEC-*Acly*^−/−^ mice that developed 100% mortality within two weeks, no CEC-*Acly*^−/−^;*Trp53*^−/−^ mice died (**Fig. S4D**). In fact, ABX failed to disrupt the colon mucosal structure (**Fig. S4E**), failed to induce SASP secretion (**Fig. S4F**), and failed to induce colonic senescent markers (**Fig. S4B**) in CEC-*Acly*^−/−^;*Trp53*^−/−^ mice. These observations demonstrate acetyl-CoA deficiency triggers p53-dependent colonic senescence that disrupts mucosal homeostasis and induces severe systemic inflammation.

### Targeted depletion of senescent cells rescues mice from developing local and systemic inflammation

The induction of p16^Ink4a^ is a universal hallmark of senescent cells. The p16-3MR transgenic mice express a p16 promoter-driven tri-modality reporter (3MR) that contains the functional domains of luciferase (Luc), RFP and ganciclovir (GCV)-sensitive HSV thymidine kinase^41^ (**Fig. 3A**). The mice have been used to quantify, visualize and eliminate senescent cells in several disease models^41–43^. To confirm that colonic senescent cells are the cause of local and systemic inflammation, we generated CEC-*Acly*^−/−^;p16-3MR mice (**Fig. 3A**). Following ABX, CEC-*Acly*^−/−^;p16-3MR mice showed time-dependent increases in SA-β-gal staining (**Fig. 3B**), luciferase activity (**Fig. 3C**) and RFP-positive cells (**Fig. 3D**) in the colonic mucosa. Importantly, treatment of CEC-*Acly*^−/−^;p16-3MR mice with GCV completely prevented ABX-induced premature death (**Fig. 3E**) and normalized the colonic epithelial structure (**Fig. 3F**). Consistently, GCV treatment blocked the induction of mucosal luciferase (**Fig. 3G**), the increase in mucosal RFP-positive cells (**Fig. 3H**) as well as the elevation of circulating inflammatory cytokines (**Fig. 3I**) in ABX-treated CEC-*Acly*^−/−^;p16-3MR mice. Western blot analysis confirmed that GCV treatment dramatically attenuated the induction of the senescence markers in the colonic mucosa of ABX-treated CEC-*Acly*^−/−^;p16-3MR mice (**Fig. 3J**). These data provide compelling evidence demonstrating that colonic senescence is the cause of systemic inflammation that leads to premature death of CEC-*Acly*^−/−^ mice.

**Figure 3.**
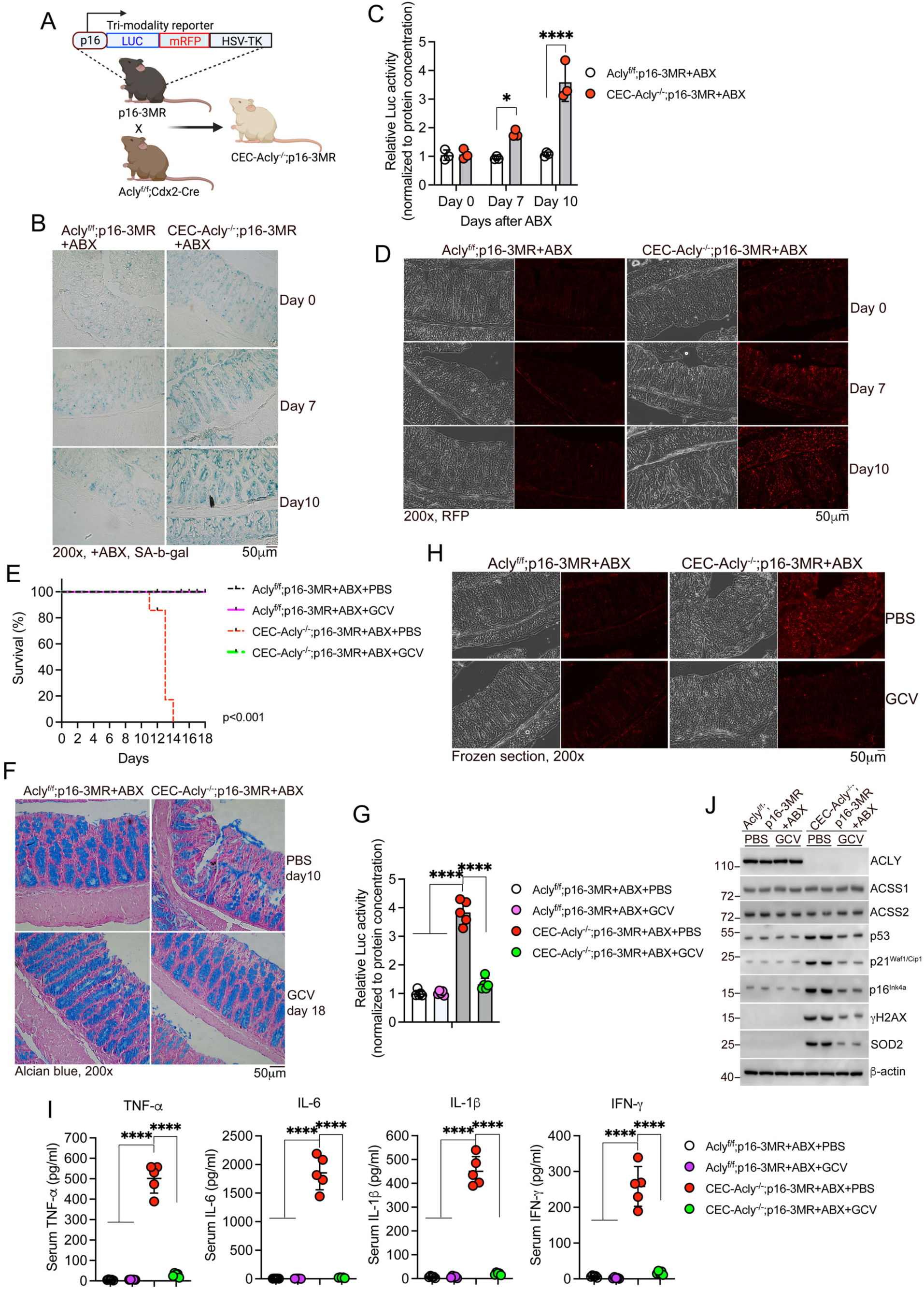
Targeted depletion of senescent cells from colonic mucosa rescues *Acly*-ablated mice from development of local and systemic inflammation and premature death. (A) Schematic illustration of the tri-modality reporter in p16-3MR transgenic mice and the generation of CEC-Acly^−/−^;p16-3MR mice; (B) SA-β-gal staining in frozen colon sections from *Acly*^f/f^;p16-3MR and CEC-*Acly*^−/−^;p16-3MR mice on days 0, 7 and 10 post ABX; (C) Luciferase (Luc) activity in purified colonic crypts from *Acly*^f/f^;p16-3MR and CEC-*Acly*^−/−^;p16-3MR mice on days 0, 7 and 10 post ABX; (D) RFP-positive cells in colonic mucosa from *Acly*^f/f^;p16-3MR and CEC-*Acly*^−/−^;p16-3MR mice on days 0, 7 and 10 post ABX; (E) Kaplan-Meier survival curves of *Acly*^f/f^;p16-3MR and CEC-*Acly*^−/−^;p16-3MR mice following ABX and ganciclovir (GCV) treatment; (F) Alcian blue stained colon sections of *Acly*^f/f^;p16-3MR and CEC-*Acly*^−/−^;p16-3MR mice on day 10 after ABX and PBS treatment, and on day 18 after ABX and GCV treatment; (G) Luciferase (Luc) activity in purified colonic crypt lysates from *Acly*^f/f^;p16-3MR and CEC-*Acly*^−/−^;p16-3MR mice on day 10 post ABX and PBS or GCV treatment; (H) RFP-positive cells in colonic mucosa from *Acly*^f/f^;p16-3MR and CEC-*Acly*^−/−^;p16-3MR mice on day 10 post ABX and PBS or GCV treatment; (I) Serum cytokine concentrations in *Acly*^f/f^;p16-3MR and CEC-*Acly*^−/−^;p16-3MR mice on day 10 post ABX and PBS or GCV treatment; (J) Western blot assessment of senescence hallmarks in colonic crypt cells from *Acly*^f/f^;p16-3MR and CEC-*Acly*^−/−^;p16-3MR mice on day 10 post ABX and PBS or GCV treatment. Data are presented as mean ± SD. *p<0.05; **p<0.01; ***p<0.001, ****p<0.0001 by two-way ANOVA.

### ACSS2 catalyzes acetate conversion to acetyl-CoA to prevent colonic senescence

Given that acetate supplementation rescued ABX-treated CEC-*Acly*^−/−^ mice, we further assessed the role of microbe-derived acetate by transplanting CEC-*Acly*^−/−^ mice with acetate-producing bacteria (**Fig. S5A**). In this experiment, we used *B. pseudocatenulatum* strain DFI 7.61, which secreted a high level of acetate but no other SCFAs (**Fig. S5B**). Following three days of ABX, no fecal SCFAs were detectable in CEC-*Acly*^−/−^mice (**Fig. S5C,D**), and all mice died within about two weeks as expected when being gavaged with PBS (**Fig. S5E**); however, transplanted with this acetate-producing strain raised fecal acetate levels (**Fig. S5D**), rescued the CEC-*Acly*^−/−^ recipients from ABX-induced death (**Fig. S5E**), normalized their colon mucosal histology (**Fig. S5F**) and blocked the induction of senescent markers in the colon (**Fig. S5G**). These observations confirm the importance of gut bacteria-produced acetate in protecting the colon from developing senescence.

Because acetate can be converted to acetyl-CoA by ACSS1 or ACSS2, we tested the hypothesis that ACSS1 or ACSS2 generates acetyl-CoA from gut microbe-derived acetate to support the *Acly*^−/−^ epithelial cells. We generated *Acly*^f/f^;Cdx2-Cre;*Acss1*^−/−^ (CEC-*Acly*^−/−^;*Acss1*^−/−^) mice (**Fig. S6A**) that carried global *Acss1* deletion and colonic epithelial *Acly* deletion simultaneously (**Fig. S6B**), and these mice exhibited no visible abnormalities including having normal colon mucosal structure (**Fig. S6C**). These results indicate that ACSS1 is not required to produce acetyl-CoA from microbe-derived acetate for the host’s survival.

We then generated *Acly^f/f^*/*Acss2^f/f^*;Cdx2-CreERT2 mice, which allowed us to simultaneously delete *Acly* and *Acss2* from colonic epithelial cells by tamoxifen (TAM) injection (double knockout, DKO) (**Fig. 4A**). Following TAM treatment all DKO mice lost weight and died within about two weeks without ABX (**Fig. 4B,C**), due to robust colonic senescence (**Fig. 4D**) that triggered severe local and systemic inflammation (**Fig. 4E,F**) and organ damage (**Fig. 4G**), and all these abnormalities could not be rescued by acetate feeding (**Fig. 4B-F**), even though acetate feeding had substantially raised fecal acetate levels (**Fig. 4H**). As expected, the DKO mice showed disrupted colonic mucosal epithelial structure (**Fig. 4I**) and robust DNA damage in crypt epithelial cells (**Fig. 4J,K**), and these phenotypes could not be corrected by acetate feeding (**Fig. 4I,J,K**). Consistently, TAM treatment markedly reduced acetyl-CoA concentrations in the cytoplasm and nucleus, but not in mitochondria, in colonic epithelial cells of *Acly^f/f^*/*Acss2^f/f^*;Cdx2-CreERT2 mice, and acetate feeding had no effects on the intracellular acetyl-CoA levels (**Fig. 4L**). Therefore, simultaneous depletion of *Acly* and *Acss2* from colonic epithelial cells phenocopied intestinal bacteria-depleted CEC-*Acly*^−/−^ mice. These observations confirm that gut microbe-derived acetate contributes to the colonic epithelial acetyl-CoA pool via ACSS2 to protect the host from developing colonic senescence.

**Figure 4.**
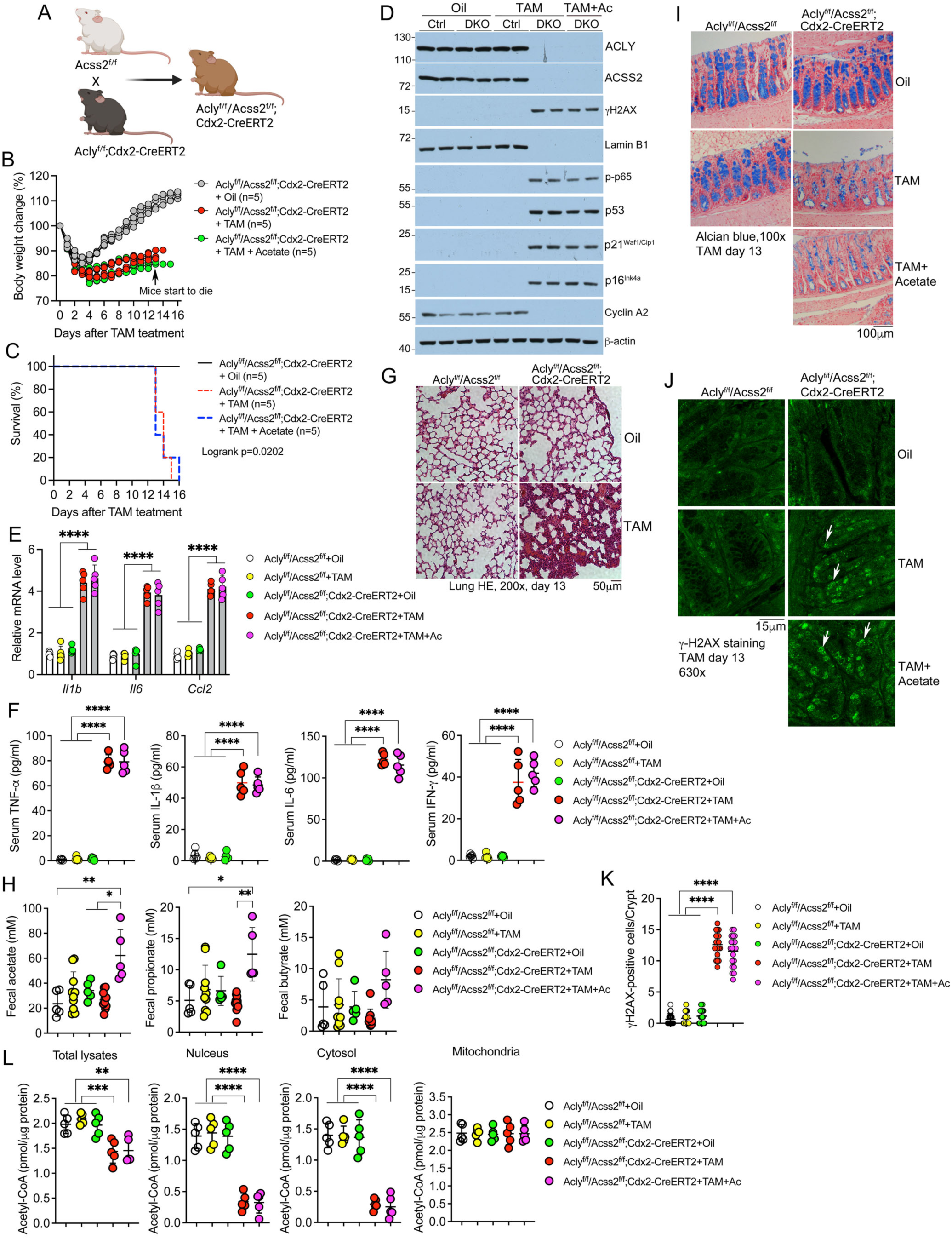
ACSS2 is required to convert microbiota-derived acetate to acetyl-CoA to prevent colonic senescence in the absence of ACLY. (A) Breeding strategy for the production of CEC-*Acly*^−/−^/*Acss*2^−/−^ double knockout (DKO) mice; (B) Body weight changes in *Acly*^f/f^/*Acss2*^f/f^;Cdx2-CreERT2 mice following tamoxifen (TAM) treatment with or without acetate (Ac) supplementation; (C) Kaplan-Meier survival curves of *Acly*^f/f^/*Acss2*^f/f^;Cdx2-CreERT2 mice following tamoxifen (TAM) treatment with or without acetate supplementation; (D) Western blot assessment of senescence hallmarks in colonic crypt cells from *Acly*^f/f^/*Acss2*^f/f^ (Ctrl) and *Acly*^f/f^/*Acss2*^f/f^;Cdx2-CreERT2 (DKO) mice on day 13 following TAM and acetate treatment; (E) RT-qPCR quantitation of SASP cytokines in colonic crypt cells from *Acly*^f/f^/*Acss2*^f/f^ and *Acly*^f/f^/*Acss2*^f/f^;Cdx2-CreERT2 mice on day 13 following TAM and acetate treatment; (F) Serum cytokine concentrations in *Acly*^f/f^/*Acss2*^f/f^ and *Acly*^f/f^/*Acss2*^f/f^;Cdx2-CreERT2 mice on day 13 following TAM and acetate treatment; (G) H&E stained lung sections from *Acly*^f/f^/*Acss2*^f/f^ and *Acly*^f/f^/*Acss2*^f/f^;Cdx2-CreERT2 mice on day 13 following TAM treatment; (H) Quantitation of fecal SCFAs (acetate, propionate, butyrate) in *Acly*^f/f^/*Acss2*^f/f^ and *Acly*^f/f^/*Acss2*^f/f^;Cdx2-CreERT2 mice on day 13 following TAM and acetate treatment; (I) Alcian blue staining of colon sections from *Acly*^f/f^/*Acss2*^f/f^ and *Acly*^f/f^/*Acss2*^f/f^;Cdx2-CreERT2 mice on day 13 following TAM and acetate treatment; (J) Confocal microscopic images of γH2AX immunofluorescence staining in colonic sections from *Acly*^f/f^/*Acss2*^f/f^ and *Acly*^f/f^/*Acss2*^f/f^;Cdx2-CreERT2 mice on day 13 following TAM and acetate treatment. *Arrows* indicate γH2AX loci in the nucleus of crypt epithelial cells; (K) Quantitation of γH2AX-positive cells in colonic mucosa of *Acly*^f/f^/*Acss2*^f/f^ and *Acly*^f/f^/*Acss2*^f/f^;Cdx2-CreERT2 mice on day 13 following TAM and acetate treatment; (L) Quantitation of acetyl-CoA concentrations in total cell lysates and nuclear, cytosolic and mitochondrial fractions of colonic crypt cells from *Acly*^f/f^/*Acss2*^f/f^ and *Acly*^f/f^/*Acss2*^f/f^;Cdx2-CreERT2 mice on day 13 following TAM and acetate treatment. Data are presented as mean ± SD. *p<0.05; **p<0.01; ***p<0.001, ****p<0.0001 by two-way ANOVA.

### *Acly* deletion triggers senescence in colonic organoids

To create a viable *Acly*-deficient organoid system, we derived colon organoids from *Acly^f/f^*;Cdx2-CreERT2 mice (**Fig. 5A**). These organoids grew normally, but once *Acly* depletion was induced by 4-hydroxytamoxifen (4-OHT) treatment, canonical senescence hallmarks including SA-ý-gal, p53, p21^Waf1/Cip1^ and p16^Ink4a^, as well as SASP (IL-6, TNF-α, and *Il1b, Il6, ccl2*), were strongly elevated in a time-dependent manner (**Fig. 5B,C,D,E**). Other senescence biomarkers including the elevation in γH2AX (DNA damage) and p-p65 (inflammation) and the decline in Lamin B1 (loss of nuclear integrity) and Cyclin A2 (cell cycle arrest) were also detected in *Acly*-deleted organoids in a time-dependent manner (**Fig. 5F**). Importantly, 4-OHT-induced senescence was blocked by acetate supplementation in the media (**Fig. 5G**). We further assessed the role of ACLY in the regulation of senescence in human colonic organoids. When *ACLY* was silenced by lentivirus-mediated *ACLY*-specific shRNA, the organoids developed a unique senescence morphology (**Fig. 5H**), accompanied by a marked induction of senescence biomarkers (**Fig. 5I**); as expected, these senescence features were rescued by acetate supplementation in the media (**Fig. 5H,I**). Therefore, these organoid models mimic the senescence phenotype of the *Acly*-ablated colonic epithelium *in vivo*, and are excellent *in vitro* systems to study acetyl-CoA deficiency-induced senescence.

**Figure 5.**
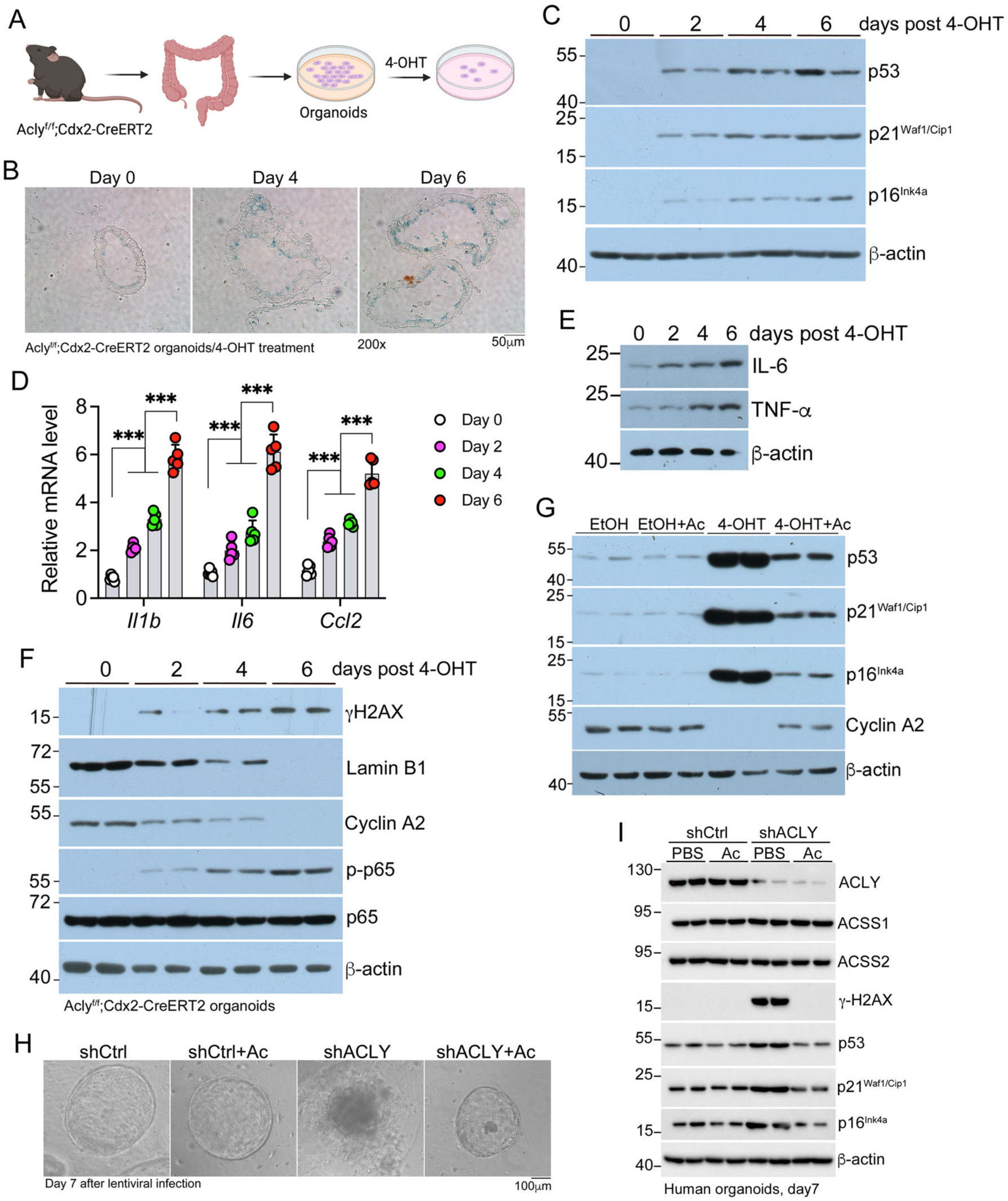
Depletion of *Acly* triggers senescence in colonic organoids. (A) Schematic illustration for the derivation of *Acly*-deleted colonic organoids from *Acly*^f/f^;Cdx2-CreERT2 mice; (B) SA-β-gal expression in *Acly*^f/f^;Cdx2-CreERT2 colonic organoids on days 0, 4 and 6 after 4-hydroxytamoxyfen (4-OHT) treatment; (C) Western blot assessment of canonical senescence hallmarks in *Acly*^f/f^;Cdx2-CreERT2 colonic organoids on days 0, 4 and 6 after 4-OHT treatment; (D) RT-qPCR quantitation of SASP cytokine expression in *Acly*^f/f^;Cdx2-CreERT2 colonic organoids on days 0, 4 and 6 after 4-OHT treatment; *p<0.05; **p<0.01; ***p<0.001, ****p<0.0001 by two-way ANOVA. (E) Western blot analysis of IL-6 and TNF-α expression in *Acly*^f/f^;Cdx2-CreERT2 colonic organoids on days 0, 4 and 6 after 4-OHT treatment; (F) Western blot analysis of senescence-related biomarkers in *Acly*^f/f^;Cdx2-CreERT2 colonic organoids on days 0, 4 and 6 after 4-OHT treatment; (G) Western blot analysis of senescence hallmarks in *Acly*^f/f^;Cdx2-CreERT2 colonic organoids treated with 4-OHT with or without acetate (Ac) supplementation in the culture media; (H) Human colonic organoids transduced with lentivirus carrying control shRNA (shCtrl) or ACLY-specific shRNA (shACLY) and cultured in the presence or absence of acetate (Ac) supplementation. Note the unique senescent morphology of shACLY-transduced organoids; (I) Western blot confirmation of ACLY knockdown, induction of senescence hallmarks, and suppression of these hallmarks by Ac supplementation in human organoids transduced with shACLY-lentivirus.

### Restoring protein acetylation rescues premature animal death and blocks colonic senescence

One main function of nucleocytosolic acetyl-CoA is to provide the substrate for protein acetylation; therefore, we compared the acetylated protein patterns in colonic epithelial cells between *Acly*^f/f^ and CEC-*Acly*^−/−^ mice by immunoprecipitation using an anti-acetyllysine antibody. There was no difference at baseline, but ABX treatment markedly reduced global protein acetylation in CEC-*Acly*^−/−^ mice, and acetate feeding reversed this change (**Fig. 6A**). Similarly, 4-OHT-induced *Acly* deletion in *Acly*^f/f^;Cdx2-CreERT2 organoids also led to a marked decrease in global protein acetylation, which was corrected by acetate supplementation in the media (**Fig. S7A**). These observations prompted a hypothesis that depletion of protein acetylation triggers colonic senescence. To test this hypothesis, we treated *Acly*^f/f^/*Acss2*^f/f^;Cdx2-CreERT2 DKO mice with Vorinostat (SAHA), a lysine deacetylase inhibitor that inhibits Class I, II and IV histone deacetylases (HDACs)^44^. Strikingly, SAHA rescued *Acly*^f/f^/*Acss2*^f/f^;Cdx2-CreERT2 mice from death following TAM-induced *Acly* and *Acss2* depletion (**Fig. 6B**), because it had mostly normalized the acetylation of colonic epithelial proteins in the DKO mice (**Fig. 6C**), and alleviated DNA damage and blocked the induction of senescence hallmarks in the DKO colonic epithelial cells (**Fig. 6D**). We confirmed that SAHA was also able to block DNA damage and senescence in 4-OHT-treated *Acly*^f/f^;Cdx2-CreERT2 organoids (**Fig. S7B**). These data indicate that the depletion of protein acetylation is a critical driver of senescence under acetyl-CoA deficiency.

**Figure 6.**
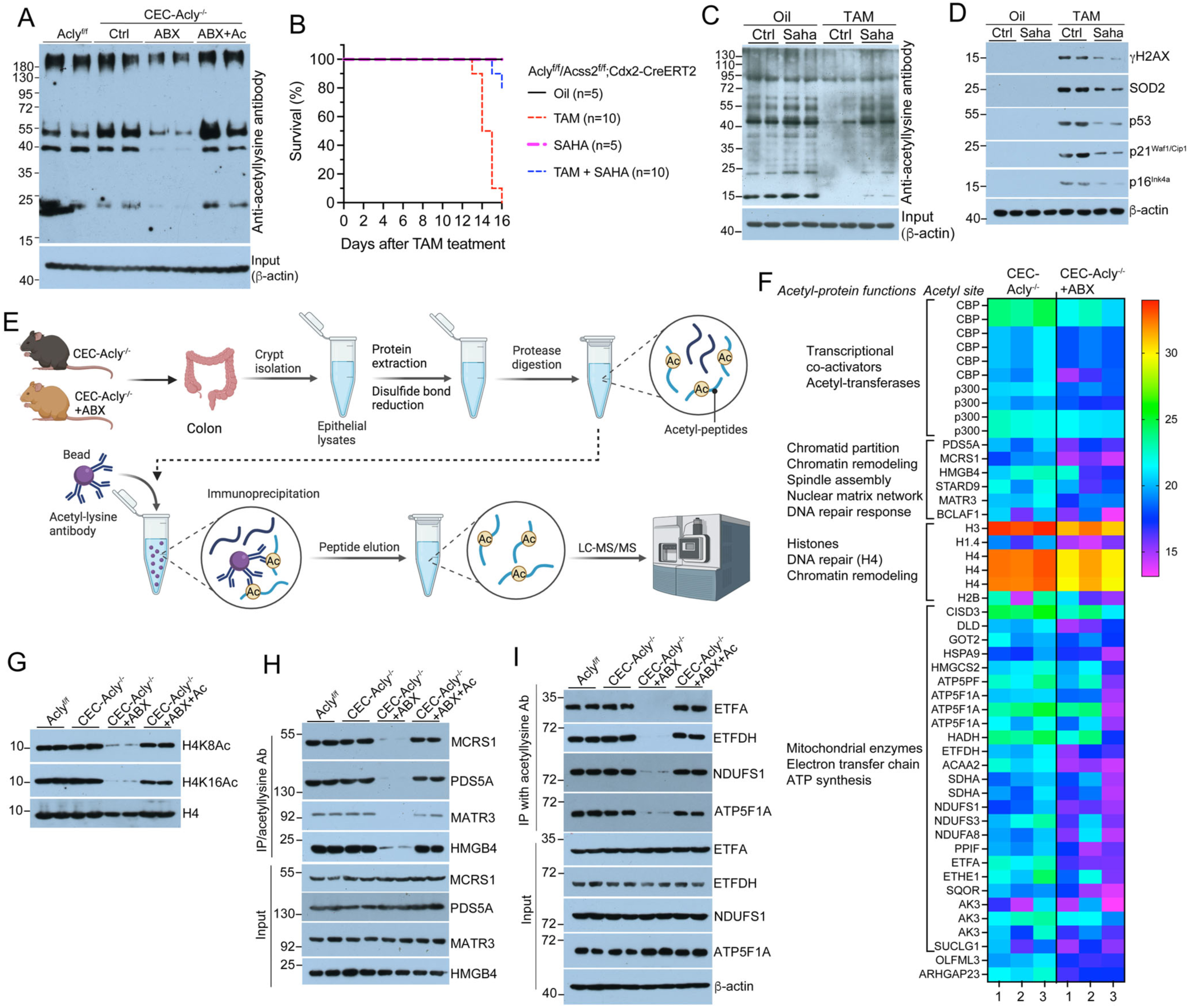
Colonic senescence is associated with the depletion of lysine acetylation in a cohort of nuclear and mitochondrial proteins. (A) Acetyl-protein profiles revealed by immunoprecipitation with anti-acetyl-lysine antibody in purified colonic crypt cells from *Acly*^f/f^ and CEC-*Acly*^−/−^ mice on day 7 following ABX and acetate (Ac) feeding; (B) Kaplan-Meier survival curves of *Acly*^f/f^/*Acss2*^f/f^;Cdx2-CreERT2 mice following tamoxifen (TAM) and SAHA treatment; (C) Acetyl-protein profiles in colonic crypt cells of *Acly*^f/f^/*Acss2*^f/f^;Cdx2-CreERT2 mice on day 13 following TAM and SAHA treatment; (D) Western blot assessment of senescence hallmarks in purified colonic crypt cells from *Acly*^f/f^/*Acss2*^f/f^;Cdx2-CreERT2 mice on day 13 following TAM and SAHA treatment; (E) Schematic illustration of acetyl-proteomics procedure; (F) Heatmap showing changes in specific lysine acetylation in a cohort of nuclear and mitochondrial proteins identified by acetyl-proteomics in CEC-*Acly*^−/−^ colonic crypt cells with or without ABX; (G) Western blot analysis of H4 acetylation at K8 and K16 in purified colonic crypt cells from *Acly*^f/f^ and CEC-*Acly*^−/−^ mice on day 7 following ABX and Ac feeding; (H) Immunoprecipitation analysis of nuclear protein acetylation using anti-acetyl-lysine antibody in purified colonic crypt cells from *Acly*^f/f^ and CEC-*Acly*^−/−^ mice on day 7 following ABX and Ac feeding; (I) Immunoprecipitation analysis of mitochondrial protein acetylation using anti-acetyl-lysine antibody in purified colonic crypt cells from *Acly*^f/f^ and CEC-*Acly*^−/−^ mice on day 7 following ABX and Ac feeding.

To search for the proteins that are depleted of acetylation as a result of acetyl-CoA deficiency, we performed acetyl-proteomic assays of colonic epithelial cells prepared from CEC-*Acly*^−/−^ mice with or without ABX (**Fig. 6E**). This analysis identified a cohort of nuclear and mitochondrial proteins that showed statistically significant reduction in site-specific lysine acetylation in the ABX-treated CEC-*Acly*^−/−^cells (**Fig. 6F**). The nuclear proteins include CBP and p300 transcriptional co-activators, proteins (PDS5A, MCRS1, HMGB4, STARD9, MATR3) known to be involved in DNA repair, DNA replication, chromatin remodeling and nuclear matrix network formation^45–50^ and histones H1, H3 and H4. The mitochondrial proteins include metabolic enzymes, proteins in electron transfer chain (ETC) complexes and subunits of ATP synthase complexes. Most of these acetylation sites have not been studied in the literature. We validated, by Western blotting with H4K8/16Ac antibodies and by immunoprecipitation with anti-acetyllysine antibody, that these nuclear and mitochondrial proteins were indeed deprived of lysine acetylation in colonic epithelial cells from ABX-treated CEC-*Acly*^−/−^ mice, and the acetylation was all normalized by acetate feeding (**Fig. 6G,H,I**). Similarly, these proteins also showed depletion of lysine acetylation in TAM-treated *Acly*^f/f^/*Acss2*^f/f^;Cdx2-CreERT2 DKO mice, and the depletion was all rescued by SAHA treatment (**Fig. S7C,D,E**). These data are consistent with the notion that acetyl-CoA deficiency triggers senescence by depleting acetylation in key nuclear and mitochondrial proteins.

### Identification of acetyl-proteins and acetyl-lysine residues that regulate colonic senescence

The acetyl-proteomic data strongly suggest that cytosolic/nuclear acetyl-CoA controls site-specific lysine acetylation in critical proteins involved in DNA repair and/or mitochondrial functions to prevent colonic senescence; as such, the depletion of acetylation in these proteins caused by acetyl-CoA deficiency disrupts their activities leading to DDR and/or mitochondrial dysfunction that trigger cellular senescence. To identify these proteins, our strategy was to overexpress the acetyl-proteins in *Acly*^−/−^ organoids and ask which protein is able to rescue the organoids from developing senescence. We took this approach based on the assumption that protein overexpression can overcome the defects in its post-transcriptional or post-translational modifications^51,52^. To this end, we used a recombinant GFP-lentivirus to transduce *Acly*^f/f^;Cdx2-CreERT2 organoids to overexpress an acetyl-protein, and then assessed whether the transduced organoids develop senescence following 4-OHT treatment to delete *Acly* (**Fig. 7A**). Organoid transduction was confirmed by GFP expression, and the normal and senescent organoid morphologies were distinguished under a light microscope. As the negative and positive controls, we confirmed that empty lentivirus-transduced *Acly*^f/f^;Cdx2-CreERT2 organoids grew normally in the absence of 4-OHT, but developed a senescent morphology following 4-OHT treatment, with prominent induction of senescence hallmarks, but transduction with human *ACLY* cDNA blocked the development of 4-OHT-induced senescence (**Fig. 7B,C,D**). We found that among the acetyl-proteins tested, only *H4c1* and *Atp5f1a* cDNAs were able to rescue 4-OHT-treated organoids from developing senescence and suppress the induction of senescence hallmarks (**Fig. 7B,C,D**).

**Figure 7.**
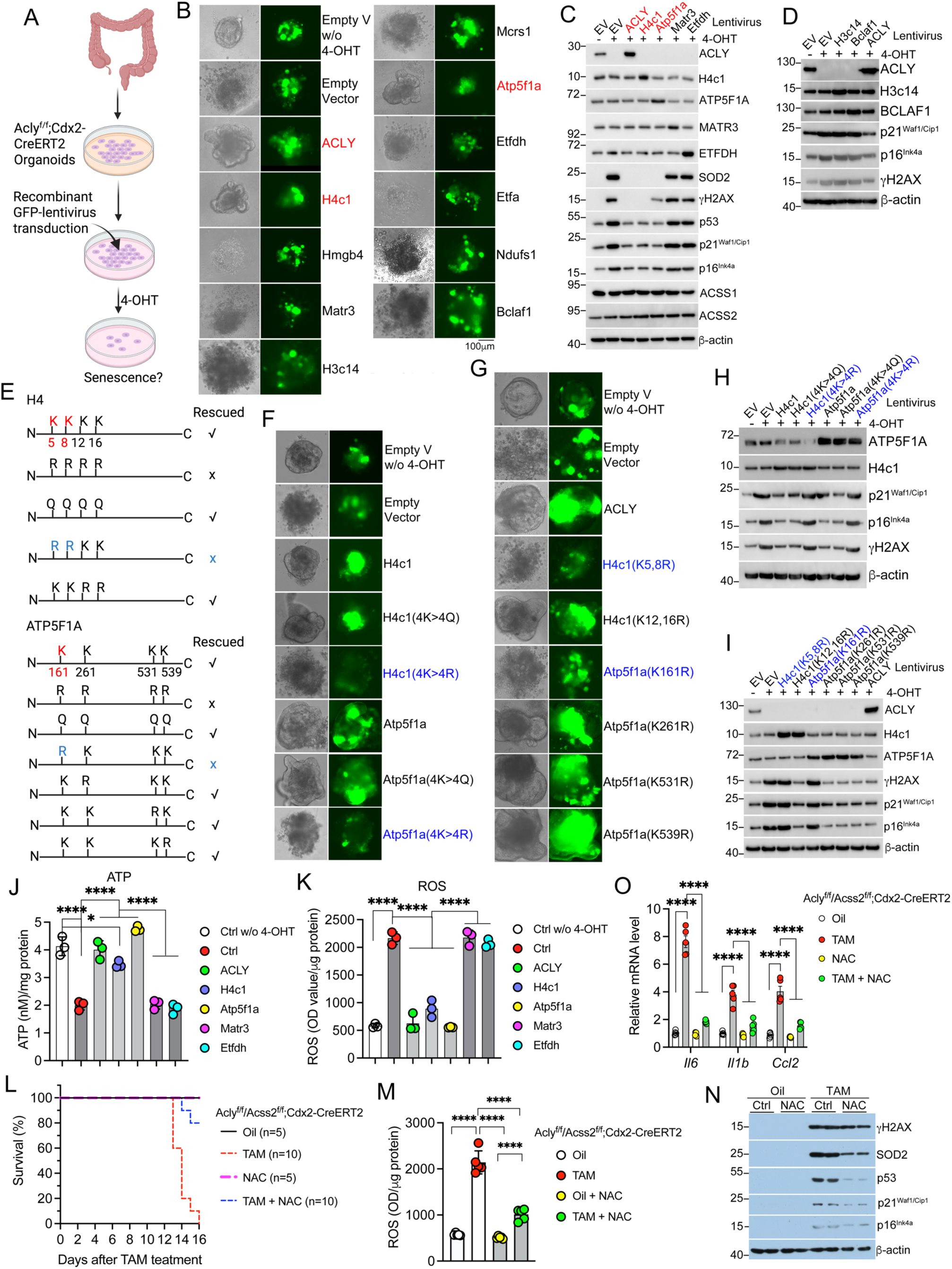
Acetylation of H4 and ATP5F1A at specific lysine residues is required to prevent acetyl-CoA deficiency-induced colonic senescence. (A) Schematic illustration of an organoid rescue strategy using recombinant lentivirus; (B) Microscopic images of *Acly*^f/f^;Cdx2-CreERT2 colonic organoids transduced with recombinant GFP-lentivirus carrying cDNA as indicated. Unless indicated, all organoids were treated with 4-OHT to deplete *Acly*. GFP fluorescence confirms the success in organoid transduction, and the normal and senescent morphology of the same organoids are identified by regular light microscopy; V, vector. (C,D) Western blot assessment of the induction or suppression of senescence hallmarks in organoids transduced with a recombinant lentivirus indicated on the top. Overexpression of each corresponding protein confirms the successful transduction of the organoids; EV: Empty vector. (E) Schematic illustration of acetylation of the lysine (K) residues identified by acetyl-proteomics in H4 and ATP5F1A, their mutation to glutamine (Q) or arginine (R) in each construct, and the outcome of the rescue experiments; (F) Microscopic images of *Acly*^f/f^;Cdx2-CreERT2 colonic organoids transduced with a recombinant GFP-lentivirus carrying wildtype H4c1 or Atp5f1a cDNA, or their mutants in which all four K residues were converted to Q or R. (G) Microscopic images of *Acly*^f/f^;Cdx2-CreERT2 colonic organoids transduced with a recombinant GFP-lentivirus carrying one of the H4c1 double mutants in which two K residues were mutated to R, or carrying one of the Atp5f1a single mutants in which each K residue was individually converted to R; (H) Western blot assessment of senescence hallmarks in *Acly*^f/f^;Cdx2-CreERT2 organoids transduced with a recombinant lentivirus carrying wildtype H4c1, H4c1(4K>4Q) or H4c1(4K>4R) mutant, or carrying wildtype Atp5f1a, Atp5f1a(4K>4Q) or Atp5f1a(4K>5R) mutant; (I) Western blot assessment of senescence hallmarks in *Acly*^f/f^;Cdx2-CreERT2 organoids transduced with a recombinant lentivirus carrying one of the H4c1 double mutants or one of the Atp5f1a single mutants as indicated; (J) Quantitation of cellular ATP concentration in *Acly*^f/f^;Cdx2-CreERT2 organoids transduced with indicated recombinant GFP-lentivirus constructs; (I) Quantitation of cellular ROS production in *Acly*^f/f^;Cdx2-CreERT2 organoids transduced with indicated recombinant GFP-lentivirus constructs; (L) Kaplan-Meier survival curves of *Acly*^f/f^/*Acss2*^f/f^;Cdx2-CreERT2 mice following tamoxifen (TAM) and N-acetylcysteine (NAC) treatment; (M) Quantitation of ROS production in purified colonic crypt cells from *Acly*^f/f^/*Acss2*^f/f^;Cdx2-CreERT2 mice on day 13 following TAM and NAC treatment; (N) Western blot assessment of senescence hallmarks in purified colonic crypt cells from *Acly*^f/f^/*Acss2*^f/f^;Cdx2-CreERT2 mice on day 13 following TAM and NAC treatment; (O) RT-qPCR quantitation of SASP cytokines in purified colonic crypt cells from *Acly*^f/f^/*Acss2*^f/f^;Cdx2-CreERT2 mice on day 13 following TAM and NAC treatment.

*H4c1* encodes H4 histone and *Atp5f1a* encodes the catalytic F1 subunit α of the mitochondrial ATP synthase complex. The acetyl-proteomics data showed that H4 polypeptide was depleted of acetylation at Lysine (K) 5, 8, 12 and 16, and ATP5F1A depleted of acetylation at K161, 261, 531 and 539 in ABX-treated CEC-*Acly*^−/−^ colonic epithelial cells. To address whether lysine acetylation at these sites is required for H4 and ATP5F1A to prevent senescence, we mutated K to Glutamine (Q) or Arginine (R) at all these sites on both proteins (**Fig. 7E**). Glutamine is an effective mimic of acetylated lysine due to the similarity of charge and chemical structure, and K to Q (K>Q) mutation as mimic of lysine acetylation is widely used in *in vitro* and *in vivo* studies for histone and non-histone proteins^53–55^, whereas K>R mutation is an acetylation-dead mutation that eliminates acetyl-lysine activity^56^. As expected, both *H4c1*(4K>4Q) and *Atp5f1a*(4K>4Q) mutants were able to rescue the organoids from senescence and suppress the induction of senescence biomarkers, but *H4c1*(4K>4R) and *Atp5f1a*(4K>4R) mutants failed to do so (**Fig. 7E,F,H**). Furthermore, to identify the acetyl-lysine sites that are critical for the anti-senescence activity, we generated *H4c1*(K5,8R) and *H4c1*(K12,16R) double mutants, and *Atp5f1a*(K161R), (K261R), (K531R) and (K539R) single mutants (**Fig. 7E**). Interestingly, while *H4c1*(K5,8R) failed to rescue the *Acly*^−/−^ organoids from senescence, *H4c1*(K12,16R) was still able to do so (**Fig. 7E,G,I**), indicating that acetylation at K5 and K8 is required to maintain H4’s anti-senescence activity. On the other hand, only *Atp5f1a*(K161R) was not able to rescue the organoids, whereas the other three *Atp5f1a* mutants (K261R, K531R, K539R) could still do so, including suppressing the induction of all senescence hallmarks (**Fig. 7E,G,I**). These results demonstrate that K161 acetylation is essential for ATP5F1A to maintain its anti-senescence activity.

DDR and oxidative stress are major drivers of cellular senescence. H4 acetylation is required for chromatin opening and DNA repair, particularly DSB repair^57–59^; thus, it is not surprising that depletion of K5/8 acetylation on H4 was found to promote colonic senescence. Mitochondrial dysfunction is a major source of reactive oxygen species (ROS) that can induce DNA damage and activate the cGAS-STING pathway ^3,37,60^. We speculated that impaired ATP5F1A acetylation, particularly the depletion of K161 acetylation, disrupts ATP synthase’s activity to generate ATP from the transmembrane H^+^ gradient leading to mitochondrial dysfunction. Indeed, we found that *Acly*^f/f^;Cdx2-CreERT2 organoids exhibited disrupted mitochondrial membrane potential after 4-OHT-induced *Acly* depletion (**Fig. S7F,G**). These organoids showed a marked reduction in ATP production and a marked increase in ROS generation, both of which were corrected by *ACLY*, *H4c1* or *Atp5f1a* transduction, but not by other acetyl-proteins (**Fig. 7J,K**). Consistently, ATP concentration was also markedly decreased and ROS levels markedly elevated in colonic epithelial cells from TAM-treated *Acly*^f/f^/*Acss2*^f/f^;Cdx2-CreERT2 mice (**Fig. S7H,I**).

Finally, treatment of these DKO mice with antioxidant N-acetylcysteine (NAC) rescued them from premature death (**Fig. 7L**), mitigated their epithelial ROS production (**Fig. 7M**), blocked senescence (**Fig. 7N**), and reduced SASP (**Fig. 7O**). We further confirmed the inhibitory effect of NAC on DNA damage, NF-κB activation and senescence marker induction in 4-OHT-treated *Acly*^f/f^;Cdx2-CreERT2 organoids (**Fig. S7J**). Collectively, these data provide strong evidence demonstrating that epithelial nucleocytosolic acetyl-CoA deficiency induces DDR and oxidative stress, as a result of acetylation depletion at strategic lysine residues in H4 and ATP5F1A proteins, which together trigger colonic senescence.

## Discussion

In this study we discovered that colonic epithelial acetyl-CoA deficiency triggers age-unrelated, p53-dependent colonic senescence that causes severe mucosal and systemic inflammation leading to premature animal death. The evidence for the development of colonic senescence is very compelling, as almost all major molecular hallmarks of senescence described in the literature are detectable, including robust inductions of SA-β-gal, p53, p21^Waf1/Cip1^, p16^Ink4a^, SASP, γH2AX and NF-κB p65 phosphorylation and marked declines in Lamin B1 and Cyclin A2. Another key piece of evidence is the requirement of p53 for the development of colonic senescence, which is a well-accepted criterion to define the occurrence of senescence^3,9,37^. Animal fatality is a very dramatic phenotype for tissue senescence. It is striking that the colonic senescent cells can secret such large amounts of SASP to induce inflammatory injury in multiple organs, which reflects the widespread and robust nature of senescence in the colon. In fact, this is confirmed by the drastic increase in p16-driven RFP-positive cells under acetyl-CoA deficiency. Moreover, targeted depletion of the senescent cells by ganciclovir confirms that these cells are the source of circulating inflammatory cytokines that cause animal death. Our data suggest that the mechanism underlying the development of colonic senescence is the selective depletion of lysine acetylation in a cohort of mitochondrial and nuclear proteins, particularly the depletion of ATP5F1A-K161 acetylation and H4-K5/8 acetylation, which disrupts the activity of mitochondrial ATP synthase to generate ATP and the activity of H4 histone required for DNA damage repair, respectively. As the mitochondrial acetyl-CoA level is normal and the TCA cycle remains intact in *Acly*^−/−^ colonic epithelial cells, defective ATP synthase is expected to disrupt oxidative phosphorylation and mitochondrial membrane potential and increase ROS production^61,62^. In fact, patients with *ATP5F1A* mutations suffer early-life mortality with oxidative phosphorylation deficiency and mitochondrial DNA depletion^63,64^. On the other hand, H4 acetylation is known to be required for DNA damage repair^57,59^. Therefore, the acetylation defects in ATP5F1A and H4 lead to increased oxidative stress and heightened DNA damage response, which together trigger cellular senescence (**Fig. S7K**). Although a variety of metabolic and nutritional factors such as NAD^+^, oxygen, metals and hyperglycemia have been reported to influence cellular senescence^22^, to our knowledge, the molecular cascade initiated by acetyl-CoA deficiency leading to colonic senescence is a previously unknown consequence of metabolic dysregulation, which represents a conceptual advance in the understanding of metabolic control of intestinal senescence.

In eukaryotic cells ACLY is the only enzyme that catalyzes the conversion of citrate to acetyl-CoA in the nuclear and cytosolic compartments; however, we found that genetic depletion of *Acly* does not render colonic epithelial cells acetyl-CoA deficient, because the cells can take up acetate, probably via MCT1, that is generated by the luminal bacteria and convert it to acetyl-CoA via ACSS2. We demonstrated that only simultaneous depletion of luminal bacteria or genetic ablation of epithelial *Acss2* in CEC-*Acly*^−/−^ mice can induce nucleocytosolic acetyl-CoA deficiency to trigger colonic senescence. Under these conditions, CEC-*Acly*^−/−^ mice started to die by day 10 and all died within two weeks because of the SASP-induced cytokine storm. Given that the intestinal epithelium renews every 4-5 days, it appears that without the supply of gut microbe-derived acetate the host’s cellular acetyl-CoA reserve can only support 2-3 cycles of epithelial renewal. We provided stable isotope tracing evidence to demonstrate the quick conversion of acetate to acetyl-CoA in colonic epithelial cells, and also showed that the supplementation of only acetate, but not propionate or butyrate, is able to rescue *Acly*-ablated colonic organoids or mice from developing senescence. We further demonstrated by bacterial transplantation that the acetate produced by luminal bacteria is indeed able to prevent CEC-*Acly*^−/−^ mice from colonic senescence and premature death. Collectively, our investigation demonstrates that gut microbe-derived acetate critically contributes to the acetyl-CoA pool of colonic epithelial cells to avert colonic senescence in situations where host’s production of acetyl-CoA from citrate is defective. This finding puts bacteria-produced acetate to a unique and important position in the maintenance of colonic epithelial homeostasis.

Gut microbiota-derived SCFAs play key roles in health and disease^65^. Bacteria produce SCFAs in the colon through fermentation of dietary fibers, which primarily include acetate, propionate and butyrate. Acetate, at ∼50 mM, is the most abundant, accounting for about 60% of the luminal SCFAs^66^. The SCFAs have been reported to participate in multiple biological processes such as regulating mucosal epithelial and immune functions and serving as energy source for the host. In this regard, there is a body of literature showing butyrate functions by activating G-protein couple receptors or inhibiting histone deacetylase^25,67–69^. In comparison, however, much less is known about acetate. Given the life-threatening consequence of colonic senescence, it makes great physiological sense that acetate, the most abundant SCFA in the colon, is used to safeguard the host’s epithelial acetyl-CoA reserve to curb the development of colonic senescence. In this regard, our work has unveiled an essential physiological function of acetate in the interplays between gut microbes and the host.

Our data showed that genetic ablation of *Acly* combined with antibiotic depletion of gut bacteria, or simultaneous deletion of *Acly* and *Acss2*, leads to a remarkable reduction of acetyl-CoA concentration in the cytosolic and nuclear compartments, but not in the mitochondria. This is not surprising because ACLY and ACSS2 are cytosolic and nuclear enzymes. Acetyl-CoA is the required substrate for protein acetylation. We found that nucleocytosolic acetyl-CoA deficiency depletes lysine acetylation in a group of nuclear and mitochondrial proteins in colonic epithelial cells. Because of the high concentration of acetyl-CoA inside the mitochondria, most mitochondrial proteins are thought to be acetylated non-enzymatically^70,71^. As such, it is surprising that the mitochondrial proteins that we identified depend on the nucleocytosolic acetyl-CoA, not the mitochondrial acetyl-CoA, for their acetylation. The nuclear proteins identified by acetyl-proteomics include proteins involved in DNA repair, DNA replication and chromatin remodeling, and the mitochondrial proteins are metabolic enzymes and subunits of ETC complexes.

Defective ETC is a major cause for ROS generation^72,73^. It is reasonable to speculate that the depletion of acetylation in these proteins most likely diminishes or disrupts their activities. Given DNA repair response and oxidative stress are major drivers of cellular senescence, we originally anticipated that most of these proteins would be able to rescue *Acly*^−/−^ organoids from senescence. Surprisingly, however, only H4 and ATP5F1A are able to do so. We further pinpointed the acetylated lysine residues in these proteins (H4-K5/8 and ATP5F1A-K161) that are essential for their anti-senescence activity, indicating that these acetyl-lysine sites are required for H4’s DNA repair activity and ATP5F1A’s ATP synthesis activity, respectively. We confirmed that acetyl-CoA deficiency indeed causes mitochondrial dysfunction, impairs ATP generation and induces oxidative stress. Therefore, the disruption of H4-K5/8 acetylation and ATP5F1A-K161 acetylation is the molecular basis that drives senescence under acetyl-CoA deficiency. However, we cannot understate the importance of the other proteins in this regard, as we cannot exclude the possibility that the guiding assumption used to design the rescue experiment – protein overexpression can overcome the impact caused by defective posttranslational modifications – is not applicable to these proteins.

Aging is associated with declines in organ functions and increases in tissue pathologies and diseases. Our finding that colonic senescence causes local and systemic inflammation has broad pathological implications, supporting the notion that tissue senescence can trigger harmful chronic inflammation leading to pathological disorders. For example, intestinal senescence has been linked to colitis^74^ and a gastrointestinal cancer^8^, the major diseases in the gastrointestinal tract. Whether and how the senescence-promoting factors identified in this work, such as acetyl-CoA deficiency and dysbiosis, are involved in the development of inflammatory diseases and gastrointestinal cancers warrants future investigation.

There are several limitations in this study. We found that acetate is able to rescue colonic senescence but cannot normalize the transcriptomic profiles of *Acly*^−/−^ colonic epithelial cells, suggesting the existence of ACLY-dependent but acetyl-CoA-independent biological and molecular processes in these cells. This work has not studied these processes nor their impacts on animal physiology. Another limitation is that it is unclear how acetyl-CoA deficiency disrupts protein acetylation and what acetyl-transferase is involved, particularly for the mitochondrial proteins. CBP and p300 have lysine acetyl-transferase activity towards histones and non-histone proteins^75^, and we found that they are depleted of lysine acetylation in the auto-acetylated regulatory loop within the catalytic acetyl-transferase domain under acetyl-CoA deficiency. Whether the acetylation-depleted CBP/p300 proteins have lost acetyl-transferase activity and are responsible for the disruption of acetylation in any of the proteins identified by proteomics have not been investigated.

## Acknowledgements

This work was supported by National Institutes of Health grants R01DK138355, R01AI151162 and 5UL1TR002389, and the Duchossois Family Institute at the University of Chicago. We thank Pieter Faber for providing sequencing services. Figures 2L, 3A, 4A, 5A, 6E, 7A, and Figures S1A, S4A, S5A, S6A, S7K were created using BioRender.com.

## Author contributions

Conceptualization: Y.C.L.; Methodology and Investigation: J.D., R.S., A.B., H.S.; Formal analysis: J.D., R.S., R.A.T, L.G., Y.L., A.B., H.S., M.O., Y.C.L.; Writing – Original Draft: J.D., Y.C.L.; Writing – Review & Editing: H.S., M.O., Y.C.L.; Fund Acquisition: Y.C.L.; Resources: L.W., M.C., M.O.; Supervision: Y.C.L.

## Correspondence

Further information and requests concerning resources and reagents should be directly addressed to Yan Chun Li.

## Materials Availability

Lentiviruses, plasmid constructs and mouse lines generated in this study will be available from the corresponding author upon request.

## Data and Code Availability

RNA-seq data generated in this study are available from GEO under the accession number GSE306668.

## METHODS

### Animals and treatments

*Acly*^f/f^ mice (*Acly^tm1.1Welk^*/Mmjax) were purchased from Mutant Mouse Resource and Research Center (MMRRC) (Stock # 043555-JAX). This genetic line has been backcrossed to the C57BL/6 background. *Acss2*^f/f^ mice (C57BL/6-Acss2tm1.2 mrl) were obtained from Taconic Biosciences (Stock #10365). CDX2-Cre transgenic mice (B6.Cg-Tg(CDX2-cre)101Erf/J; Stock # 009350), CDX2-CreERT2 transgenic mice (B6.Cg-Tg(CDX2-Cre/ERT2)752Erf/J; Stock # 022390), p16-3MR transgenic mice (B6.Cg-Tg(Cdkn2a/luc/RFP/TK)1Cmps/J, Stock # 037045) and *Trp53*^−/−^ mice (B6.129S2-*Trp53^tm1Tyj^*/J; Stock # 002101) were purchased from Jackson Laboratory. *Acss1*^−/−^ mice (ID1171 B6.129-Acss1tm1) were obtained through Center for Animal Resources and Development (CARD) (ID #1171), provided by Dr. Juro Sakai^29^. *Acly*^f/f^;Cdx2-Cre, *Acly*^f/f^;Cdx2-Cre;*Trp53*^−/−^, *Acly*^f/f^;Cdx2-Cre;*Acss1*^−/−^, *Acly*^f/f^;Cdx2-Cre;p16-3MR, *Acly*^f/f^;Cdx2-CreERT2, *Acly^f/f^*/*Acss2^f/f^*;Cdx2-CreERT2 mice were obtained through breeding. All mice were housed at 25°C and maintained in a 12h/12hr light/dark cycle. Both male and female were used in experiments. To deplete gut microbiota, mice were fed drinking water containing an antibiotic cocktail (ampicillin 1g/L; vancomycin 500 mg/L; neomycin sulfate 1g/L; metronidazole 1g/L) for up to three weeks as previously reported^32,33^. To activate CreERT2 activity, mice were intraperitoneally injected with up to three doses of tamoxifen (TAM) (one dose = 0.1ml at 10 mg/ml). Injection of corn oil (Oil) served as control. Some mice were fed drinking water containing sodium acetate (200 mM), sodium propionate (100 mM) or sodium butyrate (100 mM) for up to three weeks. Some mice were treated intraperitoneally with Vorinostat (SAHA) at 200 mg/kg (dissolved in DMSO) or N-acetylcysteine (NAC) at 50 mg/kg (dissolved in water) daily for up to three weeks. To deplete senescent cells from *Acly*^f/f^;Cdx2-Cre;p16-3MR mice, the mice were treated intraperitoneally with ganciclovir (GCV) at 25 mg/kg daily for up to three weeks. All animal study protocols were approved by the Institutional Animal Care and Use Committee at the University of Chicago.

### Colonic organoids

Mouse and human colonic organoids were grown on Matrigel as reported previously^51^. Briefly, freshly harvested colon tissues were placed in a 10 cm dish containing 5 ml of cold PBS (Ca^++^ and Mg^++^ free), cut longitudinally and rinsed 3x times with ice-cold PBS. The tissues were then cut into 2 mm pieces in 15 ml of cold PBS containing 10 mM EDTA in a 50 ml conical tube using scissors, followed by 45 min rotation (45 rpm/min) in cold room. After rotation, the tissues were precipitated by centrifugation at 4°C for 5 min (290 x g) and washed with cold PBS. The pieces were resuspended in 1 ml cold PBS containing 0.1% BSA and pipetted up and down 40 times using a pre-wetted 1 ml pipettor to make the supernatant cloudy. The supernatant was passed through 70-μm filters to enrich crypts, followed by centrifugation at 290 x g for 5 minutes at 4°C. The pellet was resuspended in cold DMEM/F-12 containing 15 mM HEPES, and the number of crypts was counted. The suspension was centrifuged again at 290 x g for 5 minutes at 4°C to precipitate crypts. The crypts were resuspended in Matrigel uniformly at 500 crypts/50 µl Matrigel, placed into a pre-warmed 24-well plate and incubated at 37°C for 20 min. After the Matrigel became solidified, pre-warmed complete Mouse IntestiCult™ Organoid Growth Medium or Human IntestiCult Organoid Growth Medium (StemCell Technologies) was added and the cells were cultured at 37°C and 5% CO_2_. The medium was changed every 2 days. In some experiments, the media was supplemented with sodium acetate (5 mM), sodium propionate (5 mM) or sodium butyrate (2 mM). For passage, colonic organoids were dissociated with the Gentle Cell Dissociation Reagent (StemCell Technologies) and re-cultured under the same condition. Human colon surgical tissues were obtained from The University of Chicago Medical Center under an IRB protocol approved by the University Medical Center Institutional Review Boards.

### Quantification of short chain fatty acids (SCFAs)

SCFAs in feces, serum and culture media were quantified by GC-MS. Fecal materials were extracted using 80% methanol spiked with internal standards at a ratio of 100 mg of fecal material/mL of extraction solvent. Samples were homogenized at 4°C on a Bead Mill 24 Homogenizer (Fisher) set at 1.6 m/s with 6×30 s cycles, 5 s off per cycle. Samples were centrifuged at 20,000 x g, –10°C for 15 min. To collect the supernatant for subsequent metabolomic analysis. Serum or media samples were extracted with 4 volumes of 100% methanol spiked with internal standards. The samples were then centrifuged at 20,000 x g, –10°C for 15 min to collect the supernatant for subsequent metabolomic analysis. Metabolites were analyzed using GC-nCI-MS and PFBBr derivatization. SCFAs were derivatized as described previously^76^ with modifications. The metabolite extract (100 μL) was added to 100 μL of 100 mM borate buffer (pH 10) (ThermoFisher), 400 μL of 100 mM pentafluorobenzyl bromide (Millipore Sigma) in Acetonitrile (Fisher), and 400 μL of n-hexane (Acros Organics) in a capped mass spec autosampler vial (Microliter). Samples were heated in a thermomixer C (Eppendorf) to 65°C for 1 h while shaking at 1300 rpm. After cooling to room temperature, samples were centrifuged at 2000 x g, 4°C for 5 min to allow phase separation. The hexanes phase (100 μL) was transferred to an autosampler vial containing a glass insert and the vial was sealed. Another 100 μL of the hexanes phase was diluted with 900 μL of n-hexane in an autosampler vial. Concentrated and dilute samples were analyzed using a GC-MS (Agilent 7890A GC system, Agilent 5975C MS detector) operating in negative chemical ionization mode, using a HP-5MSUI column (30 m x 0.25 mm, 0.25 μm; Agilent Technologies 19091S-433UI), methane as the reagent gas (99.999% pure) and 1 μL split injection (1:10 split ratio). Oven ramp parameters: 1 min hold at 60°C, 25°C per min up to 300°C with a 2.5 min hold at 300°C. Inlet temperature was 280°C and transfer line was 310°C. A 10-point calibration curve was prepared with acetate (100 mM), propionate (25 mM), butyrate (12.5 mM), and succinate (50 mM), with 9 subsequent 2x serial dilutions. Data analysis was performed using MassHunter Quantitative Analysis software (version B.10, Agilent Technologies) and confirmed by comparison to authentic standards. Normalized peak areas were calculated by dividing raw peak areas of targeted analytes by averaged raw peak areas of internal standards.

### Serum LPS quantitation

Mouse serum LPS concentrations were quantified using an LPS ELISA kit (LifeSpan Biosciences) according to the manufacturer’s instructions.

### Serum cytokine quantitation

Mouse TNF-α, IL-6, IL-1β and IFN-γ concentrations in the sera were quantified using commercial ELISA kits (BioLegend) according to the manufacturer’s instructions.

### Lung myeloperoxidase (MPO) activity

Lung MPO activity was determined as described previously^77^. Briefly, lung tissues were homogenized in 50 mM potassium phosphate and 50 mM hexadecyl trimethyl ammonium bromide (HTAB), sonicated, snap frozen and thawed twice, followed by addition of 50 mM potassium phosphate containing 0.167 mg/ml O-dianisidine dihydrochloride and 0.0005% hydrogen peroxide. Absorbance was read at 460 nm using an EL800 Universal Microplate Reader (BioTek Instruments).

### Isolation of colonic crypt cells

Mouse colonic epithelial cells were purified according to previously published methods^78–80^ with modifications. Freshly collected colons were cut open longitudinally, rinsed with ice-cold PBS, and then cut into 2 mm segments. The segmented tissues were incubated in cold PBS containing 10 mM EDTA at 4°C for 45 min with rotation (45 rpm/min). After centrifugation, the segments were washed with cold PBS once, and then resuspended in 1 ml of cold PBS. The suspension was pipetted up and down 40 times using a one-ml micropipettor, and passed through a 70-μm cell strainer to enrich crypts. Then the suspension was resuspended in 5 ml of 20% Percoll and overlaid on 2.5 ml of 40% Percoll in a 15 ml Falcon tube. Percoll gradient separation was performed by centrifugation at 780 x g for 20 min at 25°C. The cells in the interface were collected for RNA extraction or protein lysate preparation.

### ROS quantification

ROS production in colonic epithelial cells and colonic organoids was determined using 2’,7’-dichlorodihydrofluorescein diacetate (H2DCFDA) (Invitrogen). Freshly prepared colonic crypt cells or organoid cells were suspended in prewarmed PBS buffer containing H2DCFDA probe (10 µM) and incubated at 37°C for 30 min. The PBS buffer with the probe was removed via centrifugation, and the loaded cells were resuspended in prewarmed growth medium and incubated for 5 min at 37°C. The fluorescence intensity of the loaded cells was determined at 488 nm using a SpectraMax iD3 microplate reader (Molecular Devices). The optical density (OD) value was normalized to protein concentration in each sample.

### Measurement of epithelial luciferase activity

Luciferase activity in colonic crypt cell lysates was measured using a commercial Bio-Glo Luciferase Assay System (Promega) according to the manufacturer’s instruction.

### Acetyl-CoA quantification

Acetyl-CoA concentration in cellular lysates and subcellular fractions was determined using a commercial PicoProbe Acetyl-CoA Assay Kit (Fluorometric) (Abcam) according to the manufacturer’s instruction.

### ATP quantification

ATP concentration in colonic crypt lysates or colonic organoid lysates was determined using a commercial ATP Assay Kit (Colorimetric/Fluorometric) (Abcam) according to the manufacturer’s instruction.

### Subcellular fractionation

Nuclear, cytosolic and mitochondrial fractions were prepared based on published methods^81^. Briefly, colonic epithelial cells or organoids were washed with cold PBS and resuspended in STM buffer (250 mM sucrose, 50 mM Tris–HCl, 5 mM MgCl_2_, protease and phosphatase inhibitor cocktails), followed by 1 min homogenization. The cells were then centrifuged for 15 min at 800 x g and 4°C to precipitate the nuclear fraction. The supernatant containing mitochondrial and cytosolic fractions was further centrifuged at 11,000 x g for 10 min. After centrifugation, the supernatant was precipitated in acetone at –20°C for 1 hour and centrifuged at 12,000 x g for 5 min to collect the cytosolic fraction, and the pellet was resuspended in STM buffer and centrifuged at 11,000 x g for 10 min to collect the mitochondrial fraction.

### Mucosal permeability assays

Mouse gut mucosal permeability was assessed using FITC-dextran as reported previously^77^. Mice were fasted for 6 hours before being orally gavaged with 4,000 Da FITC-dextran (MilliporeSigma) at 200 mg/kg. After 2.5 hours, blood was collected from the tail, and serum FITC-dextran contents were measured at 530 nm wavelength using a SpectraMax iD3 microplate reader (Molecular Devices).

### Measurement of mitochondrial membrane potentials

Mitochondrial membrane potential was assessed using the MitoProbe JC-1 Assay Kit (ThermoFisher) according to the manufacturer’s instruction. Flow cytometric analysis for JC-1-stained cells was performed in a BD LSRFortessa unit (BD Biosciences) and data analyzed by FlowJo software V10.

### Bacterial transplantation

Mouse bacterial transplantation was performed according to published methods^82,83^ with modifications. *B. pseudocatenulatum* strain DFI 7.61 was grown anaerobically in BHIS (Becton Dickinson) until late log phase (OD600 ∼0.5, ∼1×10^7^ CFU/ml) at which point the culture was mixed with 20% glycerol 1:1 and stored in 1 mL aliquots at –80°C until use. The DFI 7.61 strain was derived from the fecal sample of a healthy donor and confirmed by whole genome sequencing. Recipient mice were fed drinking water containing an antibiotic cocktail (ampicillin 1g/L; vancomycin 500 mg/L; neomycin sulfate 1g/L; metronidazole 1g/L) for 3 days, followed by feeding autoclaved drinking water. The *B. pseudocatenulatum* stocks were thawed, and the recipient mice were gavaged with 0.2 ml of *B. pseudocatenulatum* on day 5 daily for 3 consecutive days. The transplanted mice were monitored daily for up to 3 weeks. Feces were collected on days 0, 3, 10 and 20 or at the time when the mice were killed for SCFA analysis.

### Stable isotope tracing

Colons were harvested from *Acly*^f/f^ and CEC-*Acly*^−/−^ mice with or without ABX for 7 days. The colons were longitudinally cut and the luminal contents removed. After 3x wash with cold PBS the tissues were cut into small pieces (<1 mm^3^) with a razor blade and incubated in DMEM supplemented with 10% dialyzed FBS at 37°C. After 3 hours [1,2-^13^C_2_]-acetate (5 mM) was added into the media and the incubation was continued for 60 min. Then the samples were collected by centrifugation at 4°C and metabolism was quenched with ice-cold 10% trichloroacetic acid (TCA). The samples were homogenized with a handheld Omni homogenizer, sonicated in ice-cold water and vortexed for 5 minutes at 2000 rpm and 4°C using a thermomixer and centrifuged at 21,000 x g and 4°C for 20 minutes. The supernatant was enriched for acetyl-CoA using Oasis HLB-96 well plate (Waters Corporation, 30mg sorbent, Cat # WAT058951). The solid phase extraction (SPE) wells were conditioned with 1 mL of methanol and equilibrated two times with 1 mL of water before loading the 10% TCA-extracts, washed with 0.5mL of water, and eluted with 25 mM ammonium acetate in methanol (3 x 400µL). Waters positive pressure-96 processor was used to carry out the SPE. The eluent was dried down using the Genevac EZ-2.4 elite evaporator (low-boiling-point solvent and with nitrogen purge) and re-suspended in 60 µL of 25mM ammonium acetate made in water. The chromatographic separation was performed on Waters XBridge BEH C18 (2.1×150 mm, 2.5µm, part # 186006709 and detected using Orbitrap IQ-X Tribrid Mass Spectrometer (ThermoFisher Scientific) with a H-ESI probe operated in positive mode. The autosampler was 4°C. Mobile phase A (MPA) was 10mM ammonium bicarbonate pH 9.4 (adjusted with ammonium hydroxide) and mobile phase B (MPB) was 90/10 methanol/water, 10 mM ammonium bicarbonate pH 9.4. LC gradient condition at flow rate of 0.2mL/min were as follows: 0 min: 0% B, 1.5min: 0% B, 8.0 min: 75% B, 10.0 min:100% B, 14.0 min: 100% B, 14.2: 0% B, 20 min: 0% B and column temperature was 40°C. The MS data were acquired using XCalibur 4.5 and a mass range (m/z) of either 700-1200 resolution 120K, RF lens: 45%, normalized AGC target: 100%, maximum injection time: 246 ms. The retention time and MS/MS (HCD-30) were matched against the reference standards. The data analysis was performed using Tracefinder 5.1 and the natural abundance corrections were performed using the IsoCor^84^. The data were normalized with protein content.

### Untargeted metabolomics

Snap-frozen large intestine tissues (n=4 each group) were pulverized to powder on dry ice using a mortar and pestle. The samples were extracted using 4/4/2 acetonitrile/methanol/water with 0.1M formic acid (20µL solvent per mg of tissue), vortexed and neutralized with 15% ammonium bicarbonate (8.7µL for the 100 µL of solvent), sonicated for 3 minutes in ice-cold water bath, vortexed on a thermomixer for 5 minutes at 4°C and 2000 rpm, and subjected to two freeze-thaw cycles. The samples were incubated on ice for 20 minutes and then centrifuged at 21,000 x g for 20 minutes at 4°C. An equal amount of supernatant was dried down using a Genevac EZ-2.4 elite evaporator. On the day of analysis, the dried extract was re-suspended in 150 µL of 60/40 acetonitrile/water, vortexed, sonicated for 3 minutes and vortexed for 5 minutes at 4°C using an Eppendorf ThermoMixer. The pooled QC samples were generated by combining equal volumes from each sample and injected regularly throughout the analytical batch. TCA cycle metabolite separation was performed using Thermo Scientific Vanquish Horizon UHPLC system and iHILIC-(P) Classic (2.1×150 mm, 5 µm; part # 160.152.0520; HILICON AB) column at basic. MS detection was done using Orbitrap IQ-X Tribrid Mass Spectrometer (ThermoFisher Scientific) with a H-ESI probe operating in switch polarity. The mobile phase A (MPA) was 20 mM ammonium bicarbonate at pH 9.6, adjusted by ammonium hydroxide addition and MPB was acetonitrile. The column temperature, injection volume and the flow rate were 40°C, 2 µL, and 0.2mL/minute, respectively. The chromatographic gradient was 0 minutes: 85% B, 0.5 minutes: 85% B, 18 minutes: 20% B, 20 minutes: 20% B, 20.5 minutes: 85% B and 28 minutes: 85% B. MS parameters were as follows: Acquisition range of 70-1000 m/z at 60K resolution, spray voltage: 3600V for positive ionization and 2800 for negative ionization modes, sheath gas: 35, auxiliary gas: 5, sweep gas: 1, ion transfer tube temperature: 250°C, vaporizer temperature: 350°C, AGC target: 100%, and a maximum injection time of 118 ms. Data acquisition was done using the Xcalibur software (ThermoFisher Scientific) and data analysis was performed using Compound Discoverer 3.3 & Tracefinder 5.1 software (ThermoFisher Scientific). Metabolite identification was done by matching the retention time and MS/MS fragmentation to the in-house database generated using the reference standards.

### Acetyl-proteomics

Acetyl-proteomic analysis of colonic crypt cells was carried out by Creative Proteomics (Shirley, NY). Briefly, cell lysates were extracted with 4 volumes of lysis buffer (8 M urea, 1% protease inhibitor, 1% phosphatase inhibitor). For protein digestion, 2.5 mg of proteins were diluted to 500 μL with lysis buffer. Disulfide bonds were reduced by 10 mM Tris(2-carboxyethyl) phosphine at 56°C for 1 hour. Reduced cysteine residues were alkylated by 20 mM iodoacetamide in the dark at room temperature for 30 min. Alkylated protein was precipitated at –20°C by adding 6 volumes of cold acetone and harvested by centrifuging at 10,000 x g for 10 min. The resulting protein pellets were re-dissolved in 1.5 mL of 50 mM ammonium bicarbonate and then subjected to overnight digestion at 37°C with trypsin using an enzyme to substrate ratio of 1:200 (w/w). The tryptic peptides (2 mg) were dissolved in NETN buffer (100 mM NaCl, 1 mM EDTA, 50 mM Tris-HCl, 0.5% Nonidet P-40, pH 8.0), mixed with pre-washed anti-acetyl-lysine antibody-conjugated beads and incubated at 4°C for 4 h with gentle shaking. The beads were harvested by centrifuging at 1,000 x g for 1 min at 4°C, and washed 4 times with 1 ml of NETN buffer and twice with deionized water. The bound peptides were eluted with 1% trifluoroacetic acid, and then subjected to Nano LC-MS/MS analysis. Nano LC: Nanoflow UPLC: Ultimate 3000 nano UHPLC system (ThermoFisher Scientific); Nanocolumn: trapping column (PepMap C18, 100Å, 100 μm × 2 cm, 5 μm) and analytical column (PepMap C18, 100Å, 75μm × 50 cm, 2 μm). Loaded sample volume: 1 μg. Mobile phase: A: 0.1% formic acid in water; B: 0.1% formic acid in 80% acetonitrile. Total flow rate: 250 nL/min. LC linear gradient: from 2 to 8% buffer B in 3 min, from 8% to 20% buffer B in 50 min, from 20% to 40% buffer B in 26 min, then from 40% to 90% buffer B in 4 min. Mass spectrometry: The full scan was performed between 300-1,650 m/z at the resolution 60,000 at 200 m/z, and the automatic gain control target for the full scan was set to 3e6. The MS/MS scan was operated in Top 20 mode using the following settings: resolution 15,000 at 200 m/z; automatic gain control target 1e5; maximum injection time 19ms; normalized collision energy at 28%; isolation window of 1.4 Th; charge sate exclusion: unassigned, 1, > 6; dynamic exclusion 30 s. Data analysis: Raw MS files were analyzed and searched against murine protein database using MaxQuant (v1.6.2.14). The parameters were set as follows: the protein modifications were Carbamidomethylation (C), oxidation (M) (variables), acetyl (K) (variables), acetyl (N-term) (variables); the enzyme specificity was set to trypsin; the maximum missed cleavages were set to 6; the precursor ion mass tolerance was set to 10 ppm, and MS/MS tolerance was 0.6 Da.

### Lentiviral constructs

All recombinant lentiviral constructs expressing the open reading frame of human or mouse cDNAs were generated by VectorBuilder using pLV[Exp]-EGFP:T2A:Puro-CMV or pLV[Exp]-EGFP/Neo-EF1A lentiviral vector. The cDNAs cloned into these vectors include hACLY[NM_001303274.1], mH4c1[NM_178192.2], mHmgb4[NM_027036.3], mMatr3[NM_001399986.1], mH3c14[NM_178216.3], mMcrs1[NM_016766.3], mAtp5a1[NM_007505.2], mEtfdh[NM_025794.2], mEtfa[NM_145615.4], mNdufs1[NM_145518.2], and mBclaf1[NM_001025392.1]. Lentiviral constructs carrying shRNAs were generated on pLV[shRNA]-EGFP:T2A:Puro-U6 lentiviral backbone (VectorBuilder). hACLY shRNA (shACLY) target sequence is 5’-GCCTCAAGATACTATACATTT-3’. All cloned DNA fragments in the constructs were validated by Sanger DNA sequencing. All lentiviruses were produced by transfecting HEK293T packaging cells with a titer of ∼10^7-8^ pfu/ml.

### Site-directed mutagenesis

Site-directed mutagenesis in *H4c1* and *Atp5f1a* cDNAs was performed on pLV[Exp]-EGFP/Neo-EF1A>mH4c1 and pLV[Exp]-EGFP/Neo-EF1A>mAtp5f1a templates using a Q5 Site-Directed Mutagenesis Kit (New England Biolabs) according to the manufacturer’s instructions. *H4c1* mutants generated for this study included H4c1(K5Q,K8Q,K12Q,K16Q) [H4c1(4K>4Q)], H4c1(K5R,K8R,K12R,K16R) [H4c1(4K>4R)], H4c1(K5,8R) and H4c1(K12,16R), and *Atp5f1a* mutants included Atp5f1a(K161Q,K261Q,531Q,K539Q) [Atp5f1a(4K>4Q)], Atp5f1a(K161R,K261R,531R,K539R) [Atp5f1a(4K>4R)], Atp5f1a(K161R), Atp5f1a(K261R), Atp5f1a(K531R) and Atp5f1a(K539R). All mutations were validated by Sanger DNA sequencing.

### Transduction of colonic organoid and organoid rescue assays

Transduction of colonic organoids with recombinant lentivirus was carried out according to a procedure previously described^85^ with modifications. Briefly, the entire contents in the plate well (organoids, Matrigel and medium) were collected into a 15 ml conical tube using a cell scraper. After centrifugation (5 min at 290 × g), the medium and Matrigel were aspirated carefully. The organoids were then dissociated into single cells by incubating in 1 ml of Accumax solution (StemCell Technologies) at 37°C. Enzymes in Accumax were washed out using cold PBS, and single cells were precipitated by centrifugation (5 min at 290 × g). The single-cell pellets were resuspended in the IntestiCult Organoid Growth Medium (StemCell Technologies) supplemented with 2.5 μM CHIR-99021 (APExBIO). Meanwhile, 0.1 ml of cold liquid Matrigel was added into a pre-warmed 48 well plate and incubated at 37°C for 20 min for solidification.

Then the single-cell organoid suspension, recombinant GFP-lentivirus (at 10 MOI) and polybrene (6 µg/ml) were mixed and placed on the solidified Matrigel, and the mixture was incubated at 37°C overnight. On day 2 (16 h later), the medium with dead cells were removed and another 0.1 ml of ice-cold liquid Matrigel was added to cover the cells. After solidification, IntestiCult Organoid Growth Medium was added to the organoid culture. Successful transduction was confirmed by observing GFP expression in the organoids under a fluorescent microscope. Two days after being transduced by a recombinant lentivirus, *Acly*^f/f^;Cdx2-CreERT2 organoids were treated with 4-OHT at 200 nM to induce *Acly* deletion, and the organoid morphology was photographed by light and fluorescent microcopies to assess the development of senescence. Senescence hallmarks were further assessed by Western blotting.

### Histology and immunohistochemical and immunofluorescent staining

Colons were harvested immediately after mice were killed. The colons were cut longitudinally, washed with ice-cold PBS and prepared as “Swiss rolls”^86^ for fixation overnight in 4% formaldehyde made in PBS (pH 7.2) at room temperature. The colon tissues were processed, embedded in paraffin wax and cut into 4 μm sections. The sections were stained with hematoxylin and eosin (H&E) for routine structural examination. Alcian blue staining was used to examine mucus production by goblet cells using a commercial Alcian Blue (pH2.5) Stain Kit (Vector Laboratories) according to the manufacturer’s protocol. For immunohistochemical staining, sections were boiled in 10mM sodium citrate (pH 6.0) for 10-15 min for antigen retrieval before being stained with primary antibodies. After washes, the sections were continuously incubated with horseradish peroxidase (HRP)-conjugated secondary antibodies and the antigens were visualized by incubating with 3,3’-diaminobenzidine (DAB) as substrate. For immunofluorescence staining, the sections were incubated with fluorescence-conjugated secondary antibodies and observed under a confocal fluorescent microscope. To visualize GFP or RFP fluorescence, the colon Swiss rolls were embedded in OCT media (ThermoFisher), frozen on dry ice and cut to 5 μm frozen sections using a cryostat, and the slides were examined under a fluorescent microscope.

### Senescence-associated β-galactosidase (SA-β-gal) staining

SA-β-gal activity on frozen colon sections was visualized using a Senescence β-Galactosidase Staining Kit (Cell Signaling Technology) according to the manufacturer’s instruction.

### RT-PCR

Total RNAs were extracted using TRIzol reagent (ThermoFisher). First-strand cDNAs were synthesized using a PrimeScript RT reagent kit (TaKaRa) or a ReverTra Ace qPCR RT kit (TOYOBO). Real time PCR was carried out in a LightCycler 480 Instrument II real-time PCR system (Roche), using SYBR Green qPCR Master Mixes (ThermoFisher). The relative amounts of transcripts were calculated using the 2^-ΔΔCt^ formula^87^, normalized to GAPDH transcript as an internal control. PCR primers are listed in Table S1.

### Acetyl-protein immunoprecipitation

Acetyl-protein immunoprecipitation assays were performed using a Signal-Seeker Acetyl-Lysine Detection Kit (Cytoskeleton) following the manufacturer’s instruction. Briefly, freshly purified colonic crypt cells or colonic organoids were suspended in BlastR lysis buffer and homogenized. The lysates were incubated with anti-acetyl-lysine antibody-conjugated beads at 4°C for 2 hours on a rotary mixer. The beads were collected by centrifugation and washed for 3 times. Acetyl-proteins were eluded by the Elution buffer and collected using spin columns provided in the kit. Eluted acetyl-proteins were separated by SDS-PAGE. Global acetyl-protein profiles were detected by Western blotting using anti-acetyl-lysine antibody as the primary antibody. Specific acetyl-proteins of interest were detected by Western blotting using primary antibodies against each of these proteins.

### Western blotting

Tissue and cell samples were homogenized in Laemmli buffer. Protein concentration was determined using a Bio-Rad DC RC protein assay kit. Protein lysates were separated by SDS-PAGE and then electroblotted onto Immobilon-P membranes (MilliporeSigma). The membranes were blotted with primary antibodies purchased commercially, followed by incubation with horseradish peroxidase-conjugated secondary antibody. Protein bands were visualized by chemiluminescence using an ECL Western Blot Substrate Kit (ThermoFisher). Detailed Western blot procedures were described previously^88^. Primary antibodies used for Western blot analyses are listed in the Key Resources Table.

### RNA-seq

Total RNAs were extracted from colonic crypt cells prepared from *Acly*^f/f^, CEC-*Acly*^−/−^, CEC-*Acly*^−/−^+ABX and CEC-*Acly*^−/−^+ABX+Ac mice each in triplicates using TRIzol Reagent (ThermoFisher). Poly(A^+^) mRNAs were subsequently purified from 3 µg of total RNAs using a Dynabeads mRNA Purification Kit (ThermoFisher) and used for library construction. RNA-seq libraries were prepared using a SMARTer Stranded RNA-Seq Kit (TaKaRa) according to the manufacturer’s instruction as reported previously^52^. The libraries were sequenced using an Illumina HiSeq 4000 System with single end 50-bp reads. Sequencing raw data were preprocessed using trim_galore v0.6.6, and reads were mapped by STAR^89^ v2.7.9a against Gencode GRCm39 (release M27) reference genome. The gene level read counts were obtained using FeatureCount via Rsubread 2.6.1. Differential expression was analyzed using R v4.1.0 and edgeR package. P-values were adjusted for multiple testing using the false discovery rate (FDR) correction of Benjamini and Hochberg^90^. Significant genes were determined based on an FDR threshold of 5% (0.05). Analyses of Reactome pathway enrichment and GO (Gene Ontology) biological process enrichment of the differentially expressed genes (DEGs) was accomplished using R package clusterProfiler^91^.

### Statistical analysis

Data values were presented as means ± SD. Most experiments were repeated at least twice. All bioinformatic analyses were conducted using samples of biological triplicates. Statistical analyses were performed using GraphPad Prism Version 9.1.0. For two group comparisons unpaired two-tailed Student’s *t*-test was used, and for three or more group comparisons ordinary one-way or two-way analysis of variance (ANOVA) was performed. Animal survival rates were estimated by the Kaplan-Meier method and groups were analyzed by the log-rank test. P values < 0.05 were considered statistically significant.

---

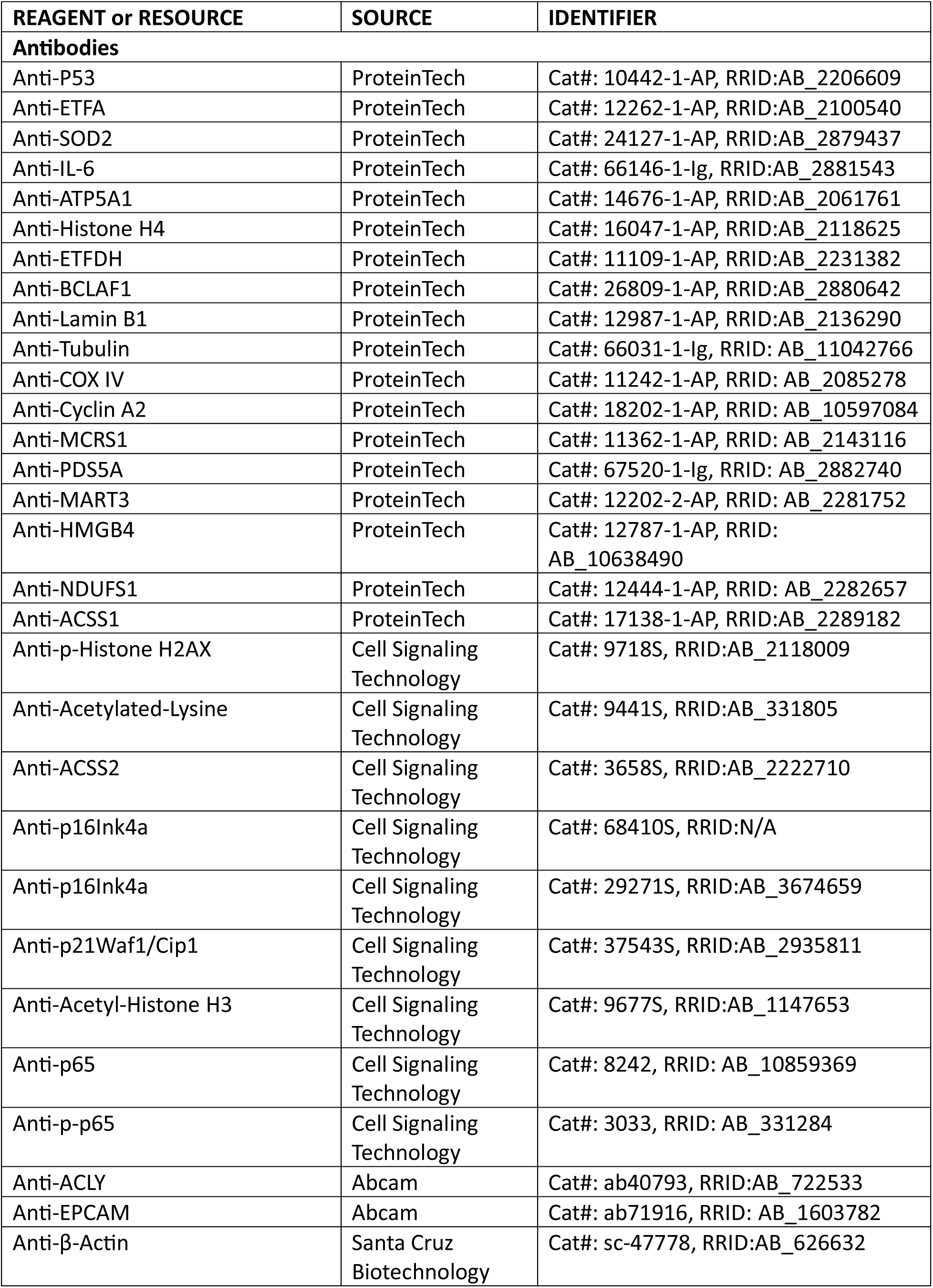
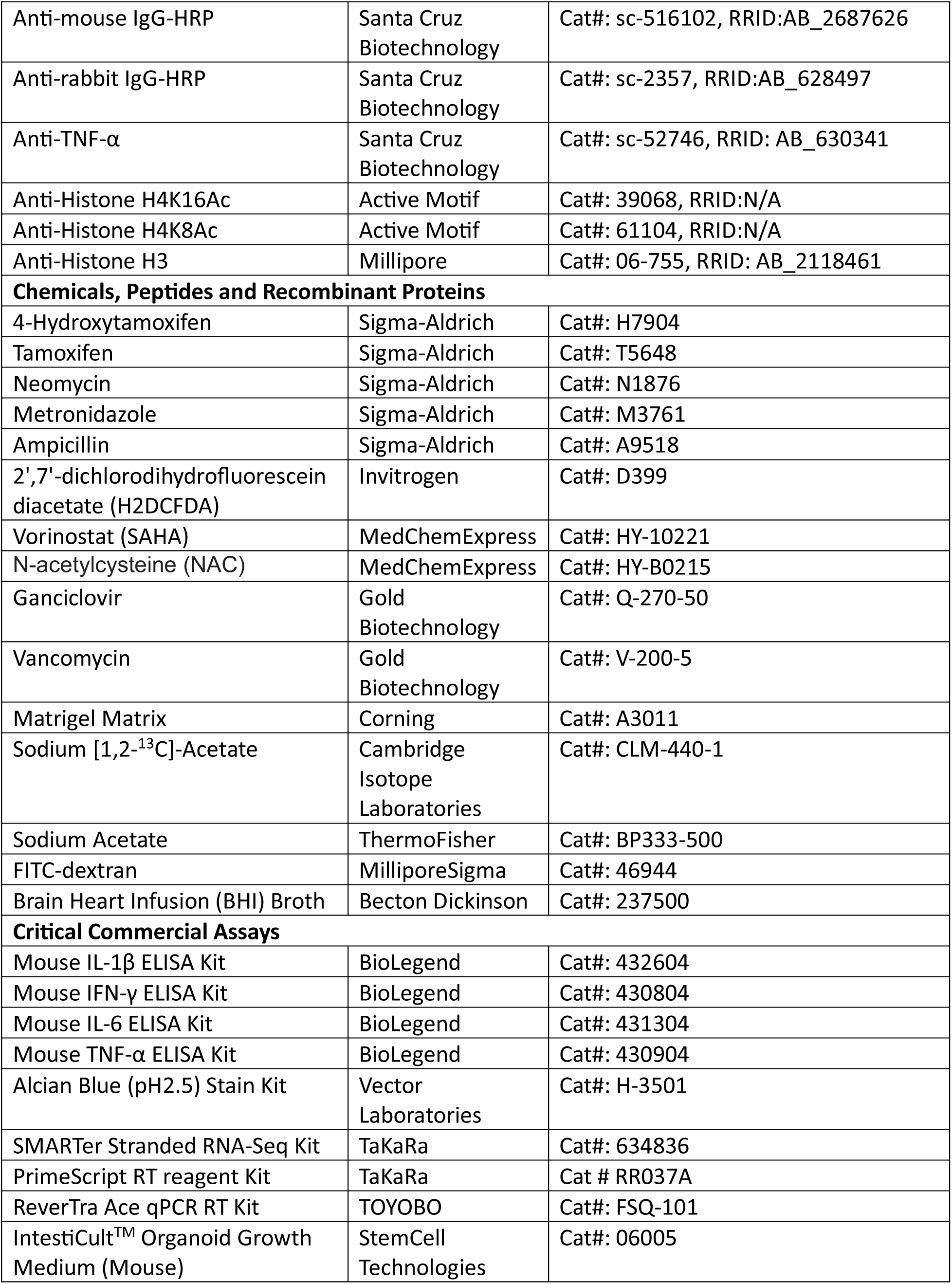
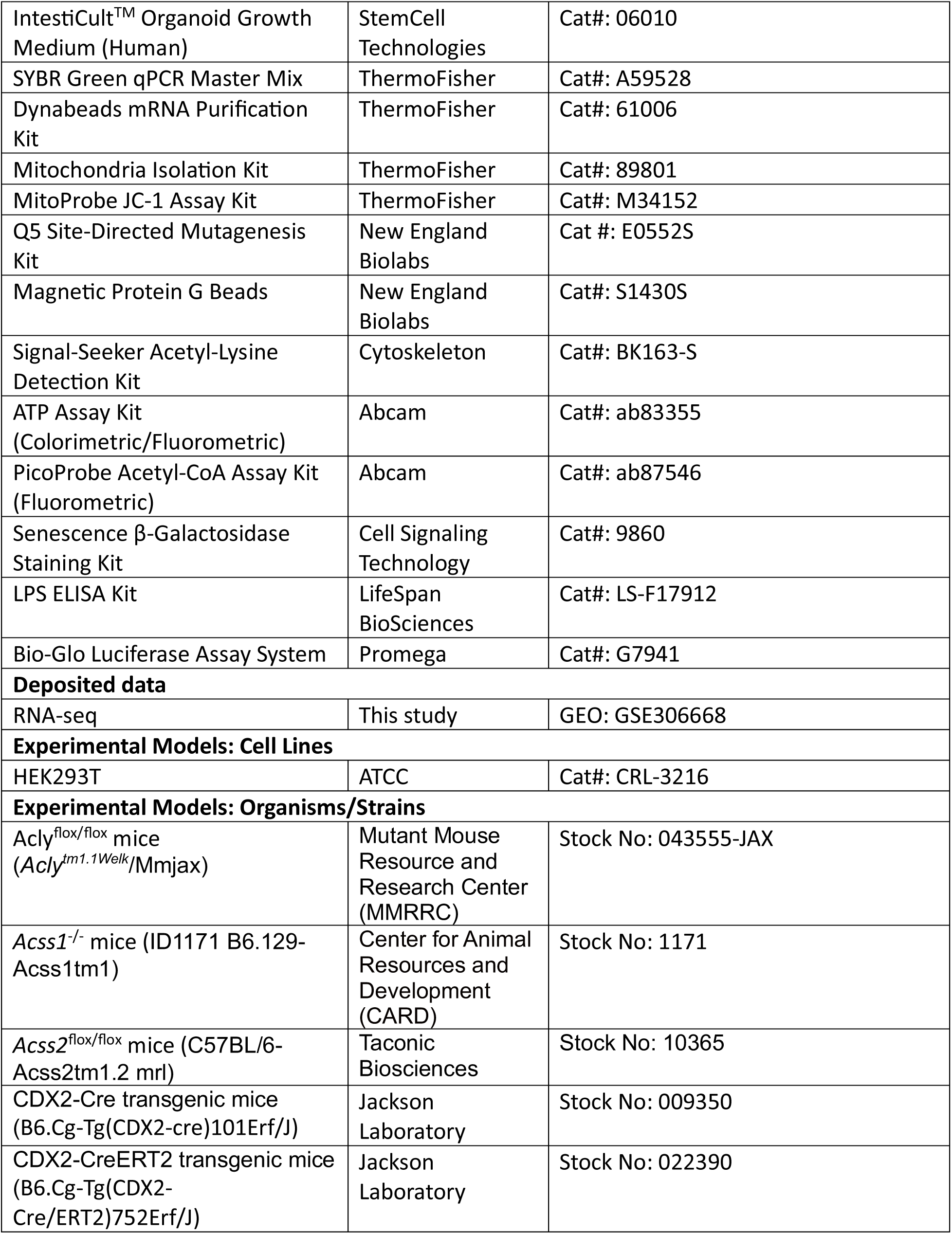
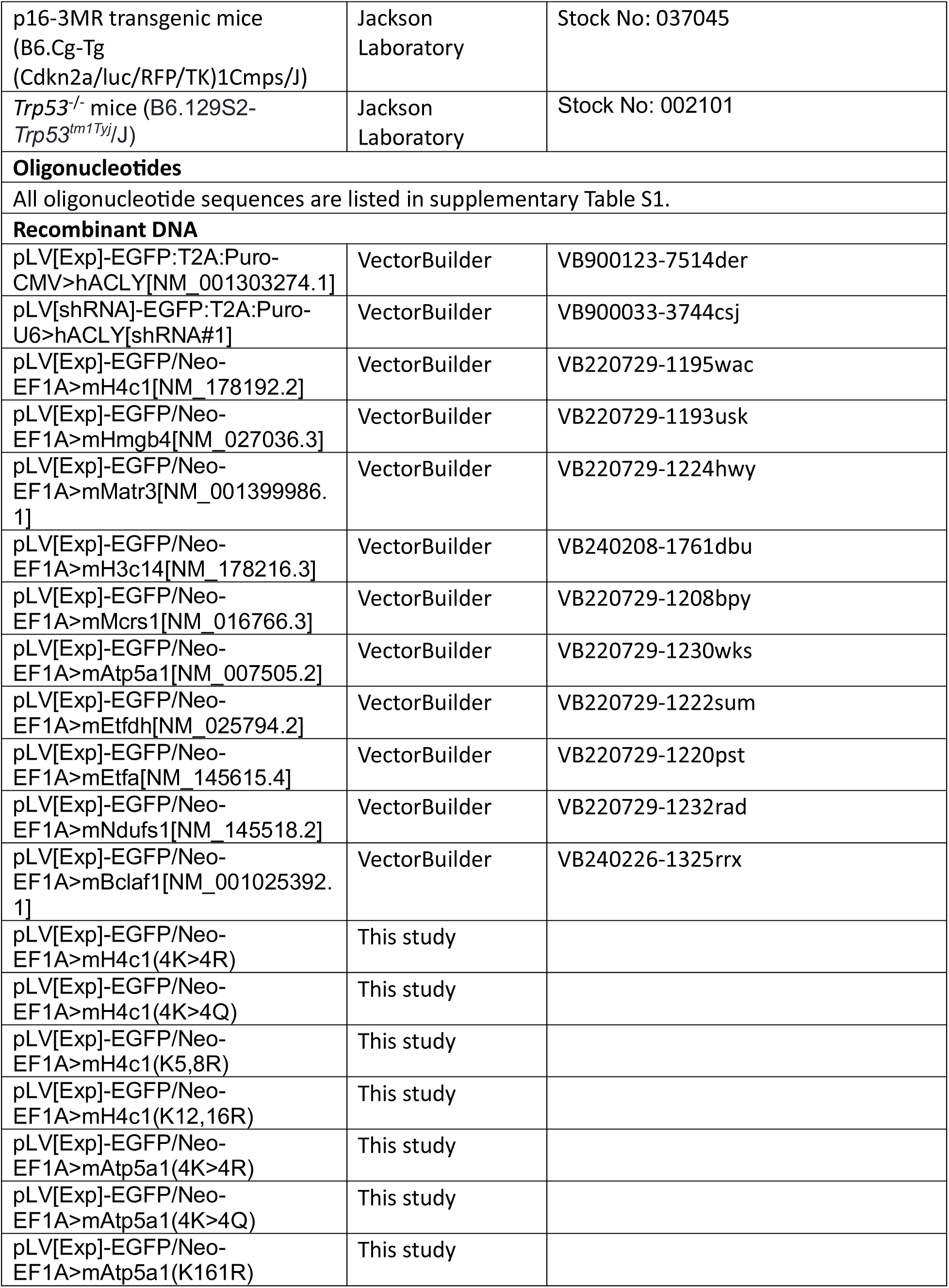
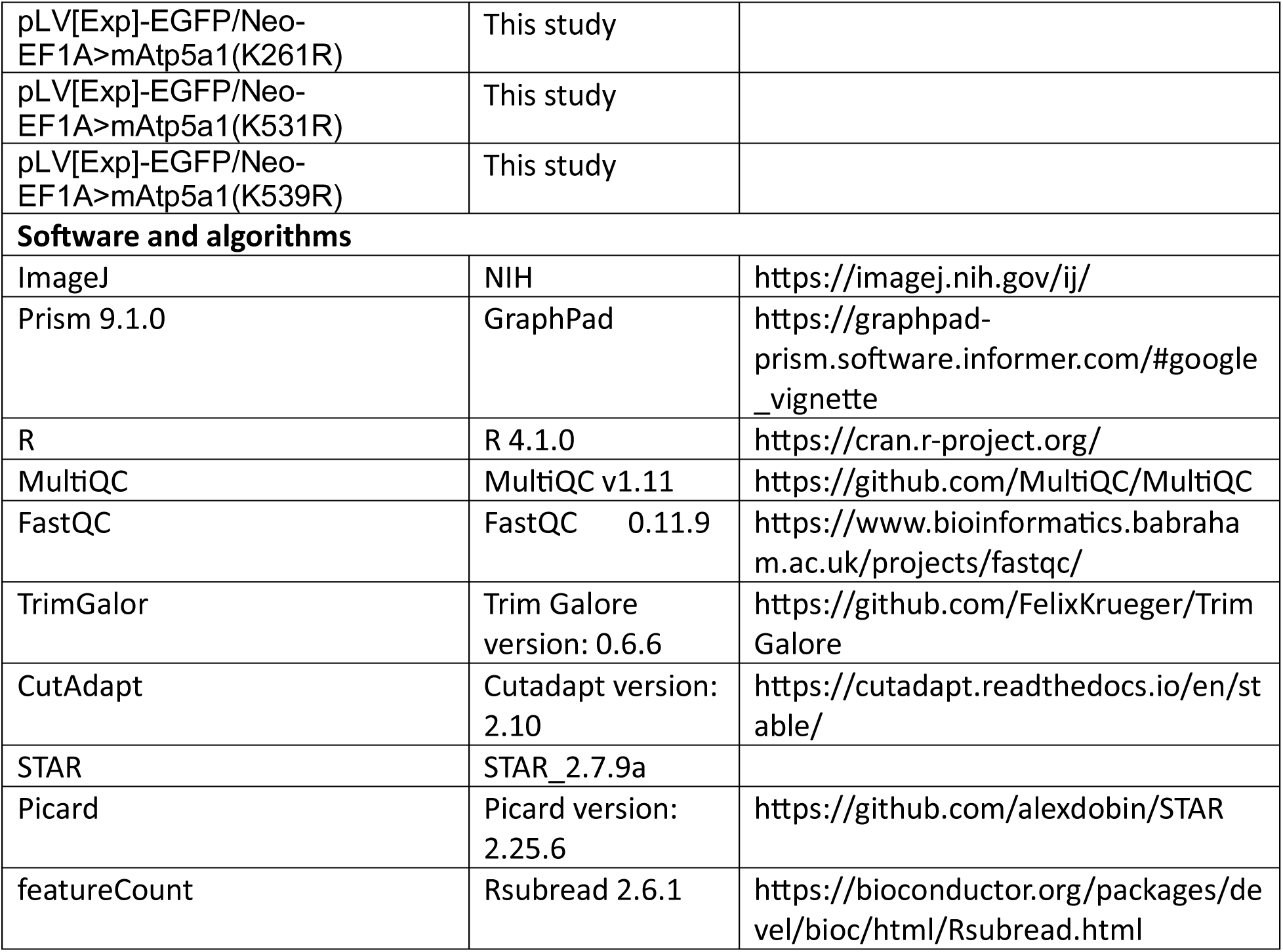

**Supplementary Table 1.**
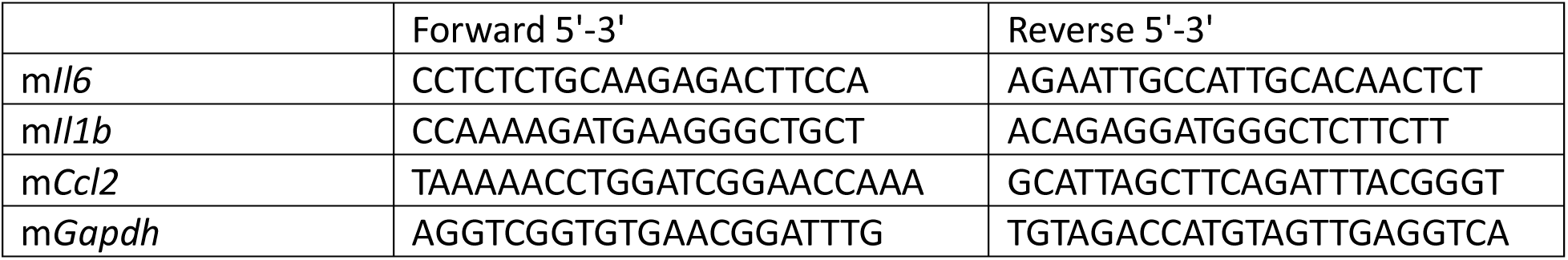
Real 2me PCR primers.

## Supplemental Figure legends

**Figure S1.**
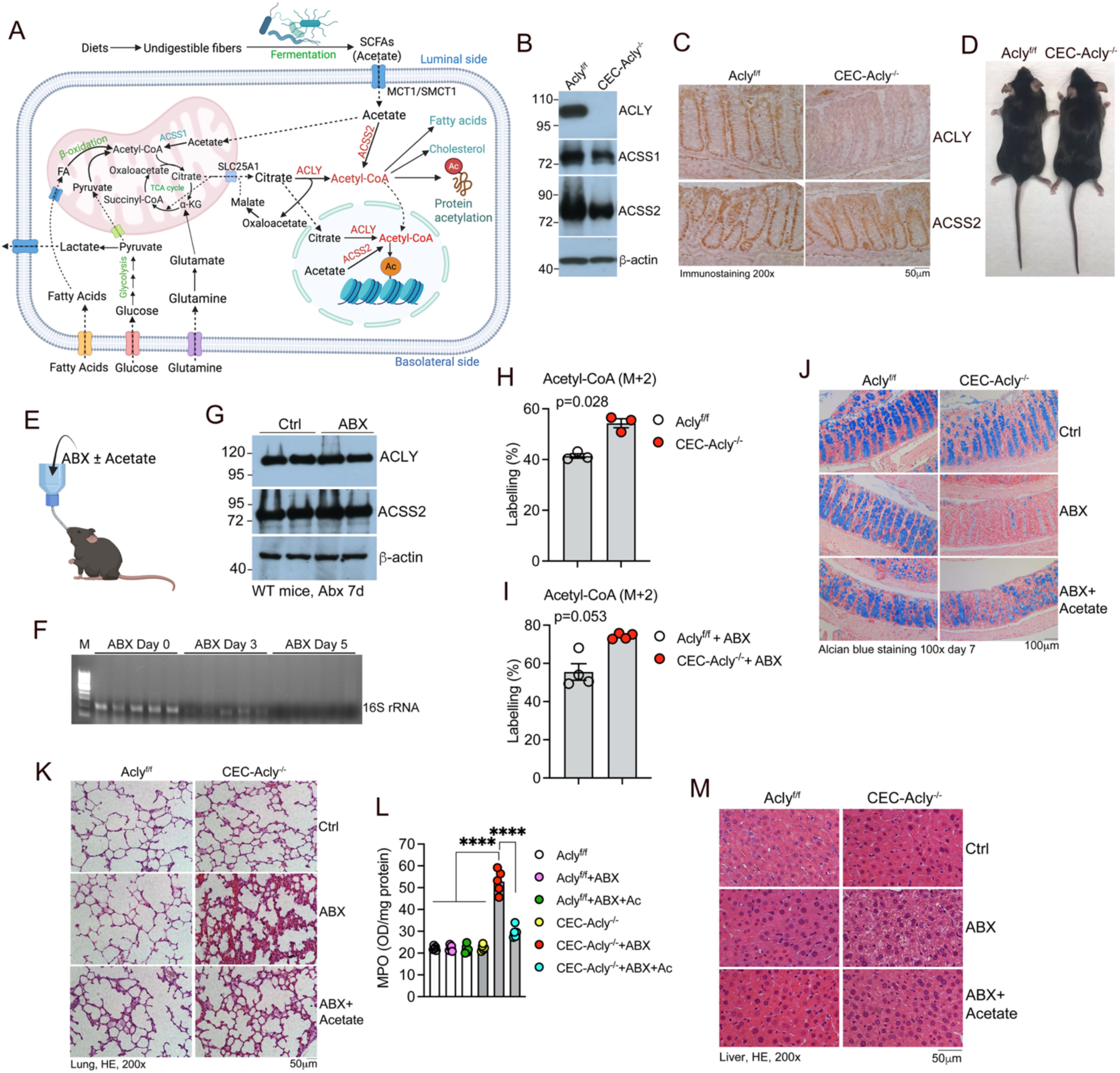
Colonic epithelial *Acly* deletion triggers systemic inflammation and multi-organ injury following antibiotic depletion of gut bacteria; related to Figure 1. (A) Schematic illustration of acetyl-CoA metabolism in colonic epithelial cells; (B) Western blot analysis of ACLY, ACSS1 and ACSS2 in purified colonic crypts from *Acly*^f/f^ and CEC-*Acly*^−/−^ mice; (C) Immunostaining of *Acly*^f/f^ and CEC-*Acly*^−/−^ colon sections using antibodies against ACLY or ACSS2; (D) Gross images of adult *Acly*^f/f^ and CEC-*Acly*^−/−^ mice; (E) Illustration of mouse treatment with an antibiotic cocktail (ABX) with or without acetate (Ac) supplementation in the drinking water; (F) PCR assessment of fecal 16S rRNA levels in mice treated with ABX for 0, 3 and 5 days; (G) Western blot analysis to assess effects of ABX on ACLY and ACSS2 expression in colonic crypt cells in wildtype mice; (H) Mass spectrometric quantitation of [^13^C]-labelled M+2 acetyl-CoA in 60 min tracing in *Acly*^f/f^ and CEC-*Acly*^−/−^ colon tissue cultures; (I) Mass spectrometric quantitation of [^13^C]-labelled M+2 acetyl-CoA in 60 min tracing in *Acly*^f/f^ and CEC-*Acly*^−/−^ colon tissue cultures from mice treated with ABX for 7 days; (J) Alcian Blue staining of distal colon sections from *Acly*^f/f^ and CEC-*Acly*^−/−^ mice on day 7 post ABX and acetate feeding; (K) H&E stained lung sections from *Acly*^f/f^ and CEC-*Acly*^−/−^ mice on day 7 post ABX and acetate feeding; (L) Quantitation of myeloperoxidase (MPO) in lung lysates prepared from *Acly*^f/f^ and CEC-*Acly*^−/−^ mice on day 7 post ABX and acetate feeding; (M) H&E stained liver sections from *Acly*^f/f^ and CEC-*Acly*^−/−^ mice on day 7 post ABX and acetate feeding.

**Figure S2.**
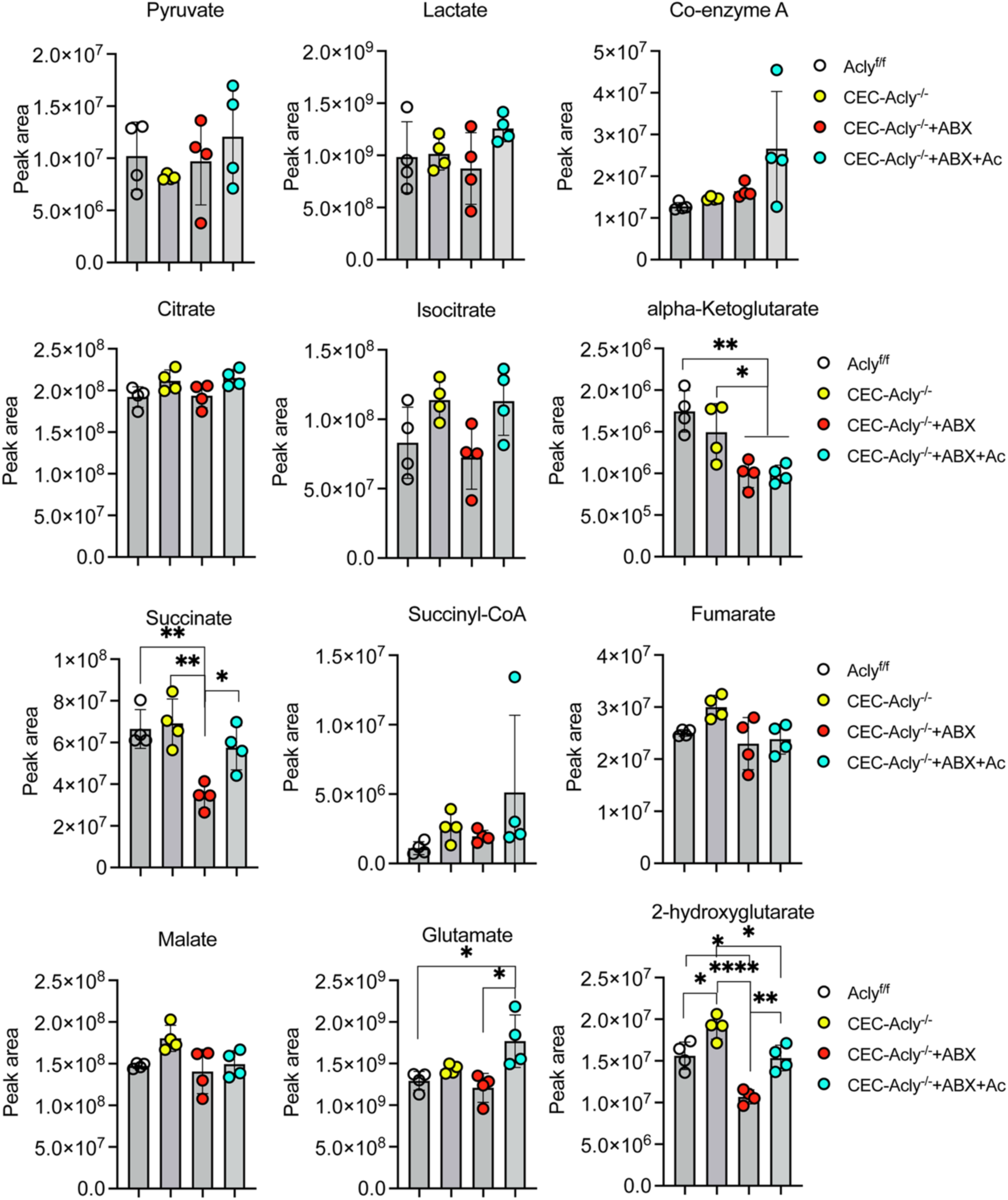
Effects of *Acly* ablation on cellular metabolites related to the tricarboxylic acid (TCA) cycle in colonic epithelial cells; related to Figure 1. Colonic crypt lysates were prepared from *Acly*^f/f^ and CEC-*Acly*^−/−^ mice on day 7 post ABX and acetate feeding, and cellular metabolites were quantified by mass spectrometry. All data are presented as mean ± SD. *p<0.05; **p<0.01; ***p<0.001, ****p<0.0001 by two-way ANOVA.

**Figure S3.**
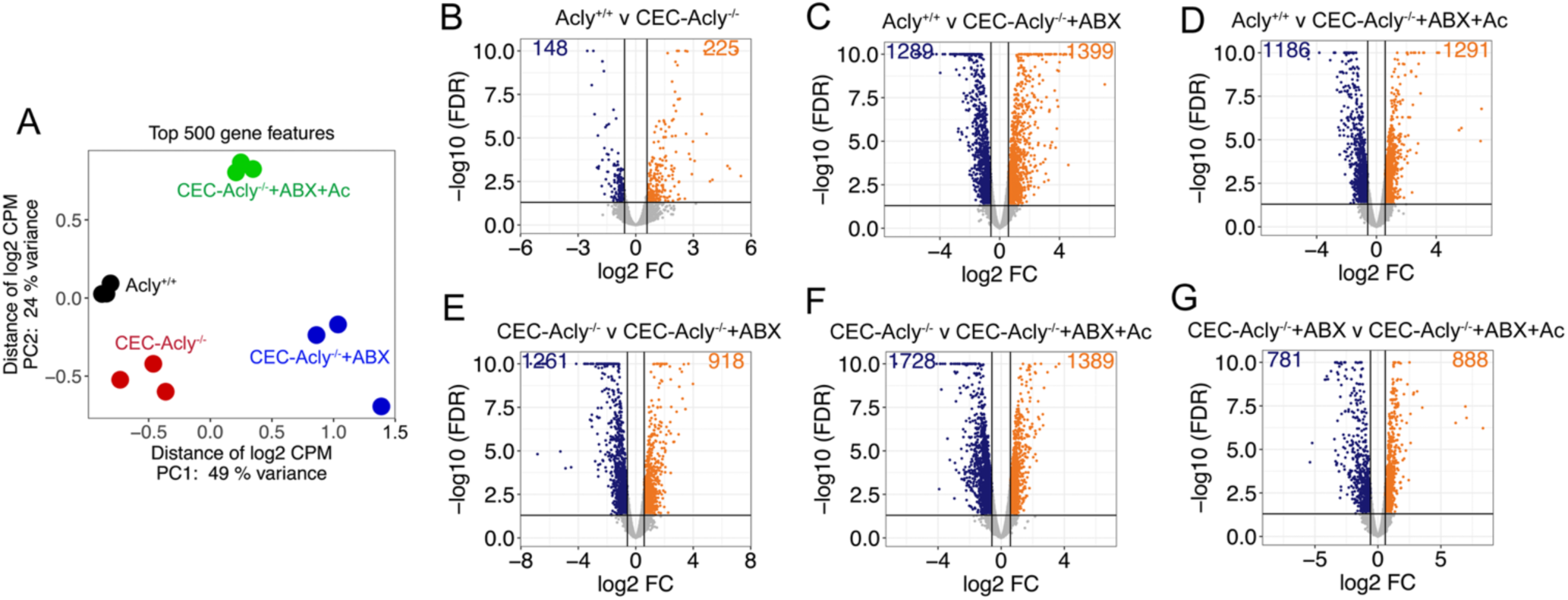
Effects of *Acly* deletion on transcriptomic profiles in colonic epithelial cells; related to Figure 2. (A) Principal component analysis of bulk RNA-seq data from *Acly*^f/f^ and CEC-*Acly*^−/−^ mice on day 5 following ABX and acetate feeding; (B-G) Volcano plots of RNA-seq data showing different comparisons as indicated. Numbers of down-regulated (*dark blue*) and up-regulated (*orange*) differentially expressed genes (DEGs) are shown on the top left and right corners of the box, respectively.

**Figure S4.**
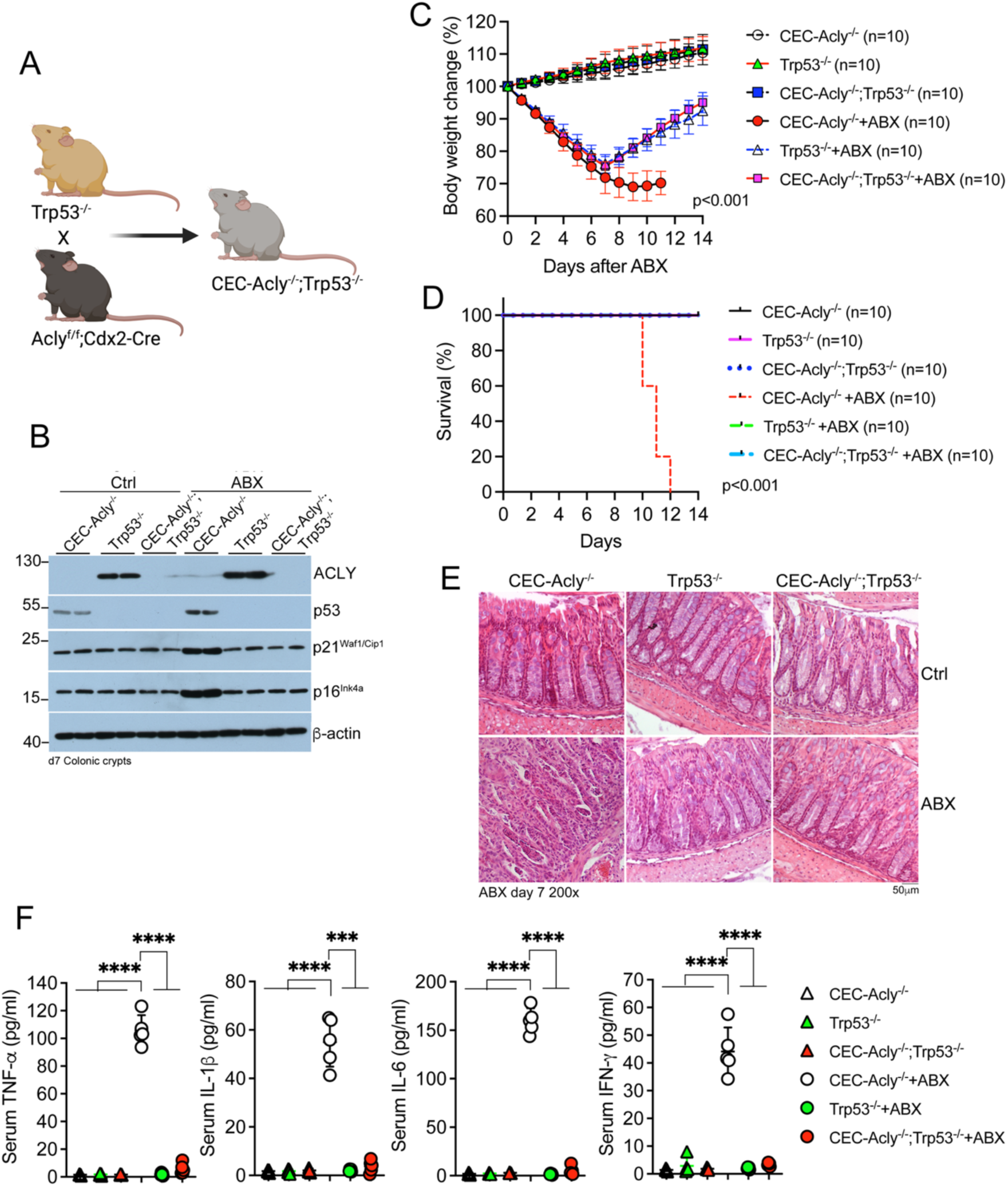
p53 is required for the development of colonic senescence and systemic inflammation triggered by epithelial acetyl-CoA deficiency; related to Figure 2. (A) Schematic illustration of the breeding strategy for the generation of CEC-*Acly^−/−^*;*Trp53*^−/−^ mice; (B) Western blot assessment of canonical senescence hallmarks in colonic crypts prepared from *Acly*^f/f^, Trp53^−/−^ and CEC-*Acly*^−/−^;*Trp53*^−/−^ mice with or without ABX for 7 days. (C) Body weight changes in CEC-*Acly^−/−^*, *Trp53*^−/−^ and CEC-*Acly^−/−^*;*Trp53*^−/−^ mice following ABX and acetate feeding; (D) Kaplan-Meier survival curves of *Acly*^f/f^, Trp53^−/−^ and CEC-*Acly*^−/−^;*Trp53*^−/−^ mice following ABX; (E) H&E stained sections of distal colons from CEC-*Acly^−/−^*, *Trp53*^−/−^ and CEC-*Acly^−/−^*;*Trp53*^−/−^ mice on day 7 following ABX and acetate feeding; (F) Quantitation of serum cytokines in CEC-*Acly^−/−^*, *Trp53*^−/−^ and CEC-*Acly^−/−^*;*Trp53*^−/−^ mice on day 10 following ABX and acetate feeding. All data are presented as mean ± SD. ***p<0.001, ****p<0.0001, by two-way ANOVA.

**Figure S5.**
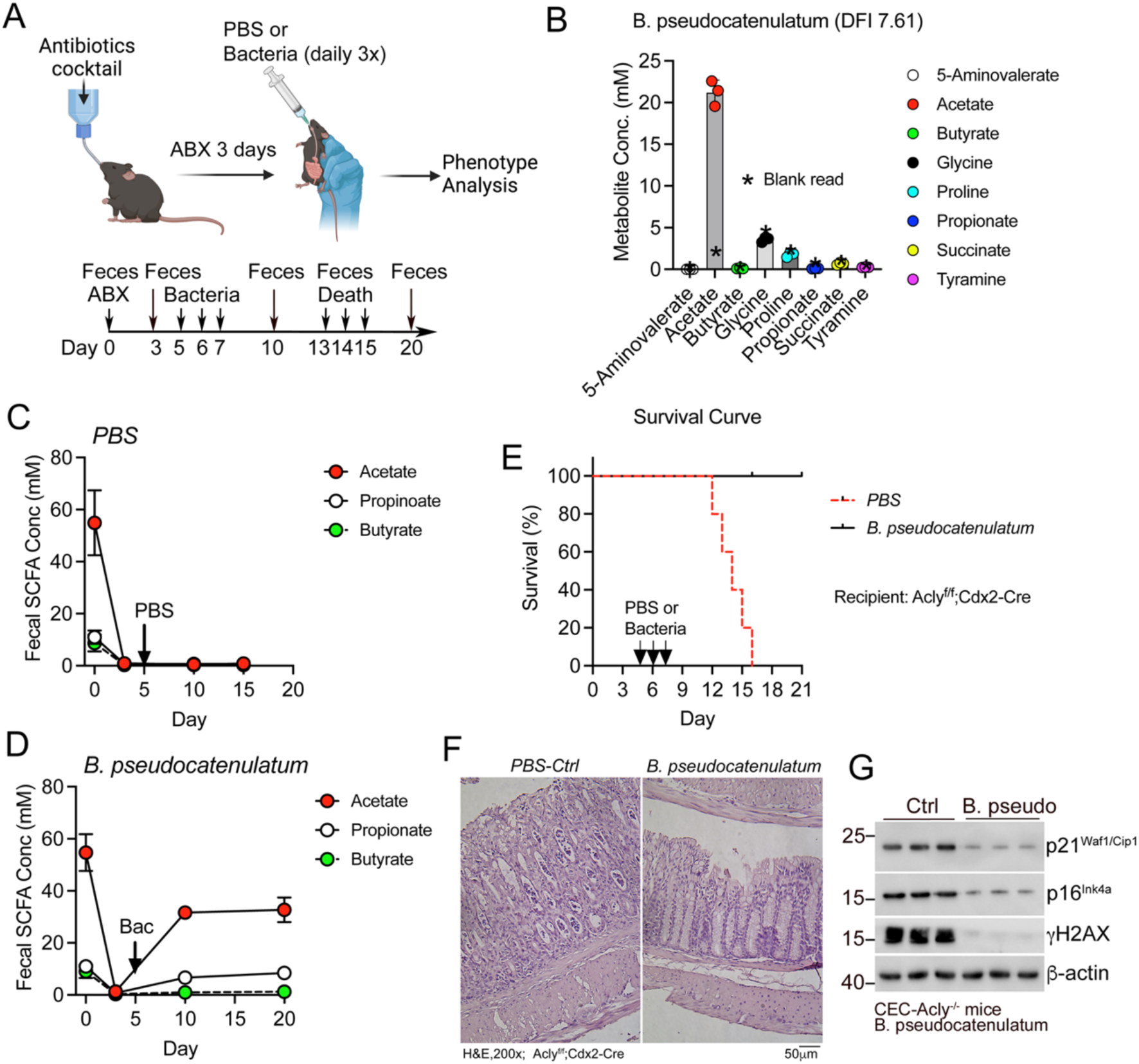
Gut bacteria-derived acetate rescues CEC-*Acly^−/−^* mice from developing colonic senescence independent of ACSS1; related to Figure 4. (A) Schematic illustration of the bacterial transplantation experiment; (B) Mass spectrometric quantitation of metabolites secreted into the media by cultured *B. pseudocatenulatum*. Blank reads (*) were obtained using culture media; (C) Time course of changes in fecal short chain fatty acids (SCFAs) in CEC-*Acly*^−/−^ recipient mice transplanted with PBS; (D) Time course of changes in fecal SCFAs in CEC-*Acly*^−/−^ recipient mice transplanted with *B. pseudocatenulatum*; (E) Kaplan-Meier survival curves of CEC-*Acly*^−/−^ recipient mice transplanted with PBS or *B. pseudocatenulatum*; (F) H&E staining of distal colon sections from CEC-*Acly*^−/−^ recipient mice transplanted with PBS or *B. pseudocatenulatum;* (G) Western blot analysis of colonic crypt cells purified from CEC-*Acly*^−/−^ mice transplanted with PBS or *B. pseudocatenulatum;*

**Figure S6.**
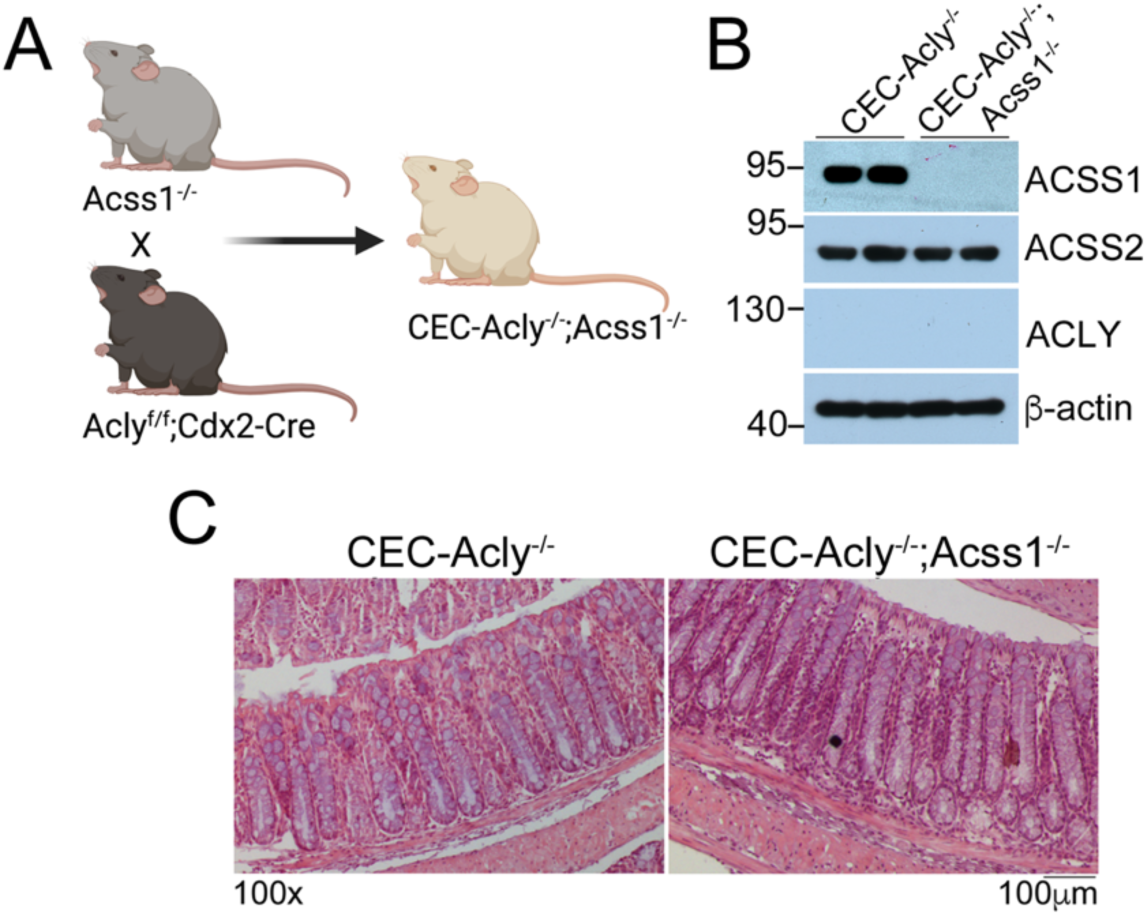
ACSS1 is not required to maintain the colonic epithelial acetyl-CoA pool; related to Figure 4. (A) Schematic illustration of the breeding strategy for the production of CEC-*Acly^−/−^*;*Acss1*^−/−^ mice; (B) Western blot analysis of ACLY, ACSS1 and ACSS2 in colonic epithelial cells isolated from CEC-*Acly^−/−^* and CEC-*Acly^−/−^*;*Acss1*^−/−^ mice; (C) H&E staining of distal colon sections from CEC-*Acly^−/−^* and CEC-*Acly^−/−^*;*Acss1*^−/−^ mice.

**Figure S7.**
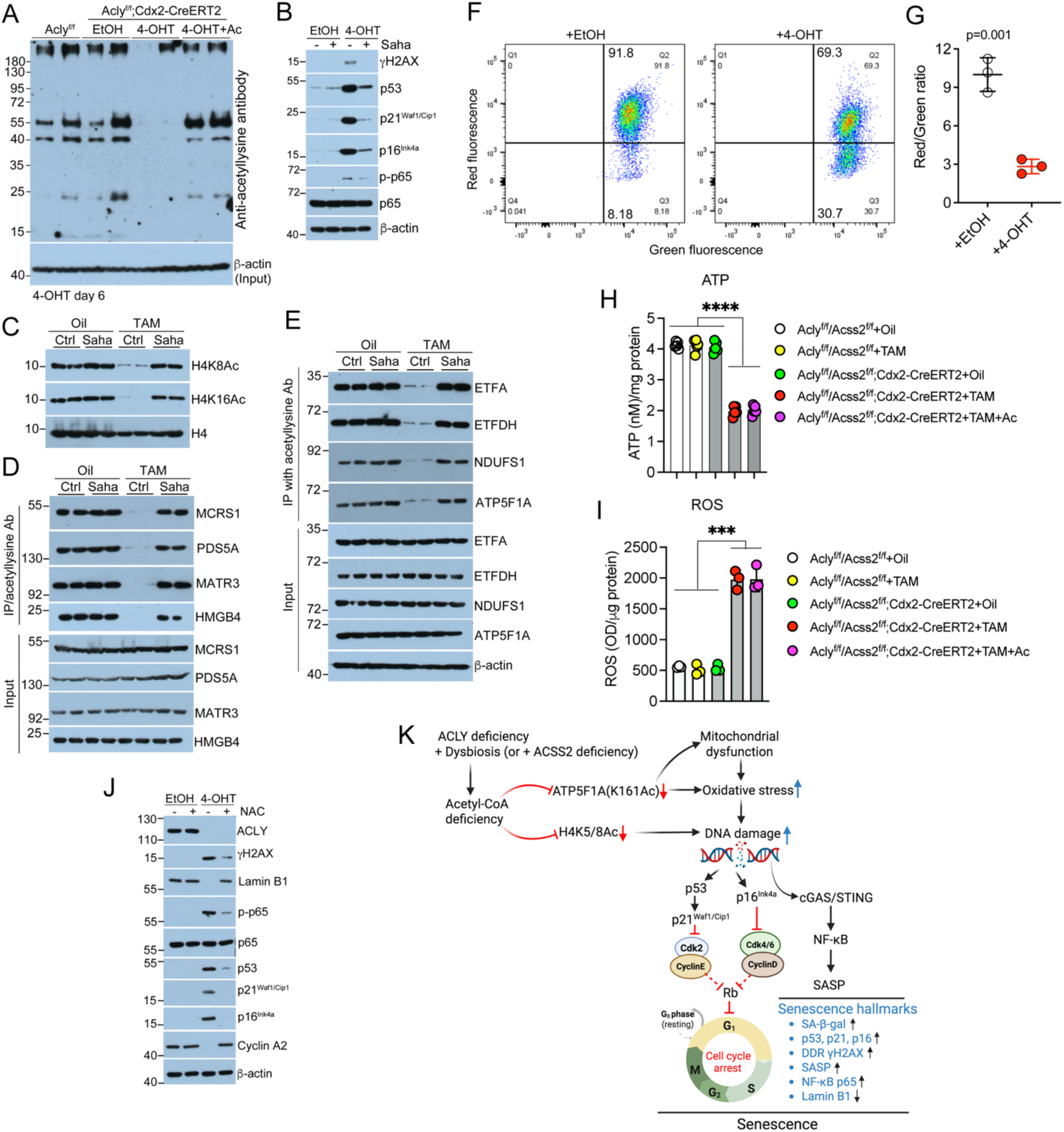
Acetyl-CoA deficiency depletes lysine acetylation in a cohort of nuclear and mitochondrial proteins leading to mitochondrial dysfunction, reduced ATP production and increased oxidative stress; related to Figures 6 and 7. (A) Acetyl-protein profiles in *Acly*^f/f^;Cdx2-CreERT2 colonic organoids on day 6 after 4-OHT treatment with or without acetate supplementation in the media; (B) Western blot analysis of senescence markers in *Acly*^f/f^;Cdx2-CreERT2 colonic organoids treated with 4-OHT and Vorinostat (SAHA); (C) Western blot analysis of H4 acetylation in colonic crypt cells from *Acly*^f/f^/*Acss2*^f/f^;Cdx2-CreERT2 mice treated with TAM and SAHA; (D) Immunoprecipitation analysis of nuclear protein acetylation using anti-acetyl-lysine antibody in colonic crypt cells prepared from *Acly*^f/f^/*Acss2*^f/f^;Cdx2-CreERT2 mice treated with TAM and SAHA; (E) Immunoprecipitation analysis of mitochondrial protein acetylation using anti-acetyl-lysine antibody in colonic crypt cells from *Acly*^f/f^/*Acss2*^f/f^;Cdx2-CreERT2 mice treated with TAM and SAHA; (F) FACS analysis of JC-1 stained *Acly*^f/f^;Cdx2-CreERT2 colonic organoids on day 6 after 4-OHT treatment; (G) Mitochondrial membrane potentials in *Acly*^f/f^;Cdx2-CreERT2 colonic organoids on day 6 after 4-OHT treatment, expressed as red (Q2) to green (Q3) ratio based on the FACS data; (H) Cellular ATP concentration in purified colonic crypt cells from *Acly*^f/f^/*Acss2*^f/f^ and *Acly*^f/f^/*Acss2*^f/f^;Cdx2-CreERT2 mice on day 13 after TAM and acetate treatment; (I) Cellular ROS production in purified colonic crypt cells from *Acly*^f/f^/*Acss2*^f/f^ and *Acly*^f/f^/*Acss2*^f/f^;Cdx2-CreERT2 mice on day 13 after TAM and acetate treatment; (J) Western blot assessment of senescence markers in *Acly*^f/f^;Cdx2-CreERT2 colonic organoids on day 6 after 4-OHT and NAC treatment. (K) Molecular mechanism whereby acetyl-CoA deficiency triggers colonic senescence. Colonic epithelial *Acly* ablation combined with intestinal bacterial depletion or *Acss2* ablation causes epithelial acetyl-CoA deficiency, which depletes lysine acetylation in a cohort of nuclear and mitochondrial proteins. Particularly, the depletion of ATP5F1A acetylation at K161 disrupts ATP synthesis and the depletion of H4 acetylation at K5 and K8 disrupts its DNA repair activity. These lead to increased oxidative stress and increased DNA damage response, which together trigger robust colonic epithelial senescence. Widespread colonic senescence causes severe systemic inflammation and premature death of the mice.

